# The Brain/MINDS Marmoset Connectivity Atlas: exploring bidirectional tracing and tractography in the same stereotaxic space

**DOI:** 10.1101/2022.06.06.494999

**Authors:** Henrik Skibbe, Muhammad Febrian Rachmadi, Ken Nakae, Carlos Enrique Gutierrez, Junichi Hata, Hiromichi Tsukada, Charissa Poon, Kenji Doya, Piotr Majka, Marcello G. P. Rosa, Hideyuki Okano, Tetsuo Yamamori, Shin Ishii, Marco Reisert, Akiya Watakabe

**Affiliations:** Brain Image Analysis Unit, RIKEN Center for Brain Science, Japan; Laboratory for Molecular Analysis of Higher Brain Function, RIKEN Center for Brain Science, Japan; Neural Computation Unit, Okinawa Institute of Science and Technology Graduate University; University Medical Center Freiburg, Germany; Department of Systems Science, Kyoto University, Japan; Laboratory for Marmoset Neural Architecture, RIKEN Center for Brain Science, Japan; Department of Physiology, Keio University School of Medicine, Japan; Center for Mathematical Science and Artificial Intelligence, Chubu University, Japan; Laboratory of Neuroinformatics, Nencki Institute of Experimental Biology of Polish Academy of Sciences, Warsaw 02-093, Poland; Australian Research Council, Centre of Excellence for Integrative Brain Function, Monash University Node, Clayton, VIC 3800, Australia; Neuroscience Program, Biomedicine Discovery Institute and Department of Physiology, Monash University, Clayton, VIC 3800, Australia

## Abstract

We report on the implementation and features of the Brain/MINDS Marmoset Connectivity Atlas, BMCA, a new resource that provides access to anterograde neuronal tracer data in the prefrontal cortex of a marmoset brain. Neuronal tracers combined with fluorescence microscopy are a key technology for the systematic mapping of structural brain connectivity. We selected the prefrontal cortex for mapping due to its important role in higher brain functions. This work introduces the BMCA standard image preprocessing pipeline and tools for exploring and reviewing the data. We developed the BMCA-Explorer, which is an online image viewer designed for data exploration. Unlike other existing image explorers, it visualizes the data of different individuals in a common reference space at an unprecedented high resolution, facilitating comparative studies. To foster the integration with other marmoset brain image databases and cross-species comparisons, we added fiber tractography data from diffusion MRI, retrograde neural tracer data from the Marmoset Brain Connectivity Atlas project, and tools to map image data between marmoset and the human brain image space. This version of BMCA allows direct comparison between the results of 52 anterograde and 164 retrograde tracer injections in the cortex of the marmoset.

## Introduction

To better understand the function of the primate brain, it is essential to map its structural connectivity using microscopic imaging data. Since the mapping of the entire connectome at single synapse resolution is still impractical due to large image size and imaging limitations, neuronal tracer injections combined with high-resolution fluorescence microscopy provide the best resolution currently achievable for systematic mapping of animal brains.

One of the most extensive open tracer image databases for mammalian brain connectivity is the Allen Mouse Brain Connectivity Atlas (***Oh et al., 2014***; ***Kuan et al., 2015***), which is the current standard for collecting, processing, and publicly sharing brain connectivity data from animal models in a systematic way. However, model organisms such as rodents have limitations when it comes to understanding primate cognition, to a large extent because of differences in brain anatomy, which translate to less complex cognitive abilities (***Schaeffer et al., 2020***; ***Carlén, 2017***).

Mental and neurological disorders, including age-related dementias, pose a major challenge to modern societies, with broad implications for economic development and well-being. Therefore, it is not surprising that there is great interest in studying the structure and function of primate brains to advance our understanding of the origin, development, and treatment of such diseases.

In recent years, the marmoset has gained popularity among primate models due to its small size, high reproductive rate, and cognitive abilities (***Okano, 2021***). For example, the marmoset has become an established model for studying Parkinson’s disease (***Ando et al., 2012***), autism spectrum disorder (***Watanabe et al., 2021***), and Alzheimer’s disease (***Sato et al., 2020***). Unlike rodents, marmosets have well-developed visual and auditory cortices, which contain the same basic subdivisions as the human brain and reflect specializations for social interaction (***Solomon and Rosa, 2014***; ***Toarmino et al., 2017***; ***Schaeffer et al., 2020***), a complex of premotor and posterior parietal areas responsible for sophisticated spatial and movement planning functions (***Bakola et al., 2022***; ***Hori et al., 2021***), and the same basic subdivisions of the prefrontal cortex (PFC) as the human brain (***Burman and Rosa, 2009***; ***Reser et al., 2013***).

Here we introduce the implementation and features of the Brain/MINDS Marmoset Connectivity Atlas (BMCA), a public access resource that provides a significant new step towards the exploration of the structural basis of primate cognition. The BMCA is built on datasets collected by the Brain/MINDS project(***Watakabe et al., 2021***), which derived from TET-amplified AAV anterograde neural tracer injections into various locations in the marmoset PFC, one of the key regions that differentiate primates from other mammals. This core database contains data from 52 anterograde neural tracer injections in adult marmosets and has complementary structural MR images. Further, for 19 of the datasets, we combined a retrograde tracer with the anterograde tracer, resulting in the ability to visualize bidirectional connections.

Automated serial two-photon tomography (STPT) (***Ragan et al., 2012***) was used to acquire the serial section images of the fluorescent anterograde tracer signals. Coronal sections were taken every 50 *μm*, with an in-plane resolution of about 1.35*μm*/px, which is sufficiently high to identify individual axon structures in the imaging plane. In addition, backlit images were taken before Nissl staining from sections that were collected after two-photon tomography, to reveal features of the brain myelination. Nissl and backlit sections were imaged under brightfield microscopy. Prior to STPT acquisition, ex-vivo whole brain MR images were acquired from marmosets using the high angular resolution diffusion imaging (HARDI) technique. All images were automatically processed, and the results were integrated into the BMCA. The BMCA gives access to the datasets in a common reference image space with a high resolution of 3 × 3 × 50*μm*^3^ that shows detailed morphology of axon fibers (Figure 1). Tools and supplementary data, such as atlas annotations and dMRI measurements, are also provided in the BMCA.

**Figure 1.**
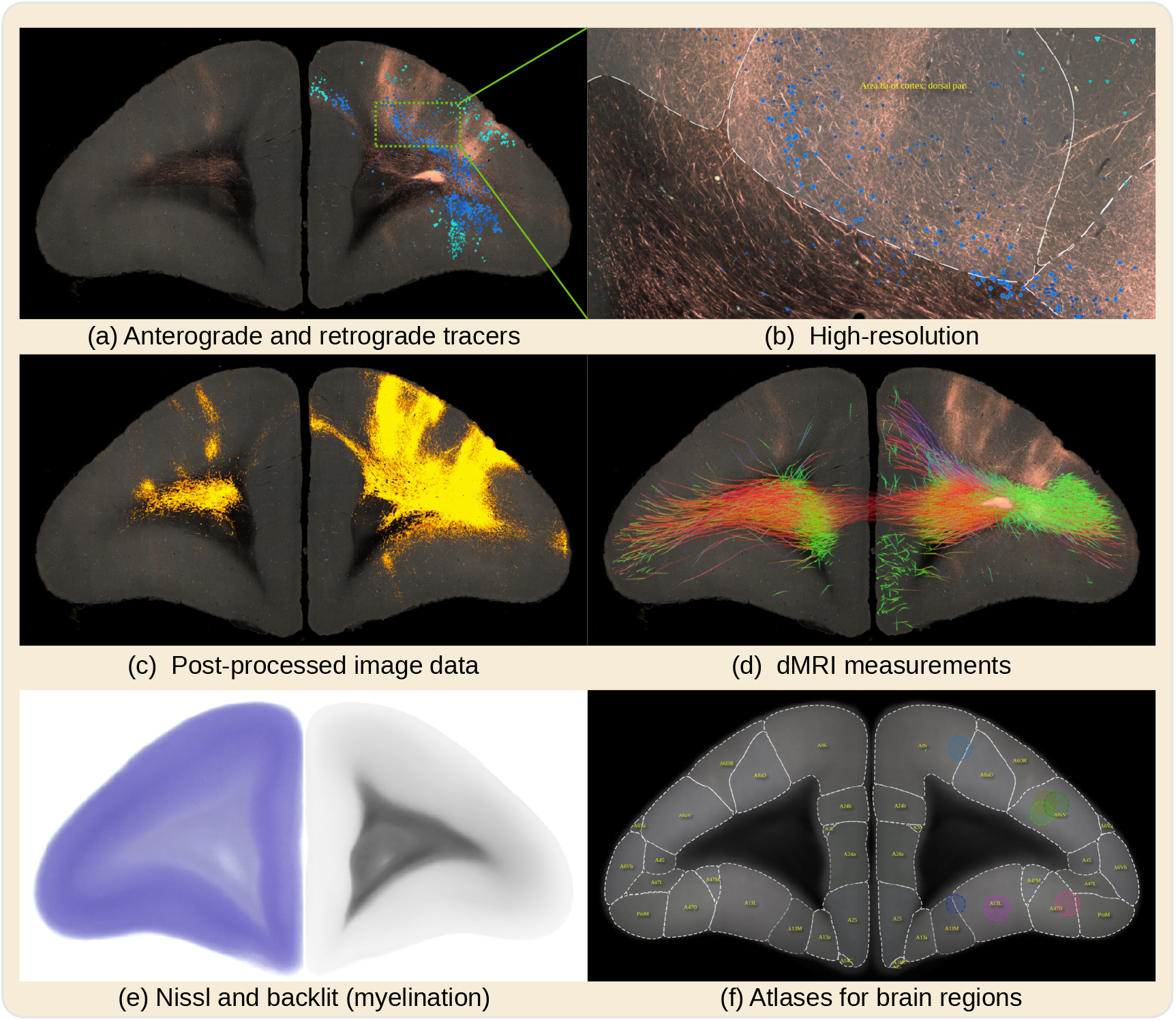
The BMCA comprises the image data of 52 anterograde tracer injections and 19 retrograde tracer injections placed into the marmoset PFC. The BMCA has been supplemented with retrograde neural tracer data from the Marmoset Brain Connectivity Atlas project (145 injections). This figure shows a series of examples of virtual coronal sections from the BMCA all in the same reference image space. Figure (a) shows a serial two-photon tomography fluorescent image of an anterograde neural tracer, and as an overlay, a retrograde tracer dataset from the Marmoset Brain Connectivity Atlas with a similar injection site (The different blue tones indicate whether cells are beneath or above layer IV). Subfigure (b) shows a detailed close-up of a portion of the same image. Figure (c) shows the segmented tracer from image (a) over the auto-fluorescent background. The overlay in subfigure (d) shows a tractography of fibers originating from the same site as the tracer injection based on the averaged dMRI data. The colors reflect fiber directions. The BMCA also includes backlit and Nissl images (see (e)) and incorporates brain region annotations from major brain atlases for marmosets (see (f) for an example of the BMA).

A surge of interest in marmoset led to the construction of various neuroinformatic resources, such as the Marmoset Brain Mapping project (MBM) (***Liu et al., 2018, 2020***), the marmoset Brain/MINDS atlas (BMA) (***Woodward et al., 2018***), and the Marmoset Brain Connectivity Atlas (MBCA) (***Majka et al., 2020***). The latter comprises a large amount of cortical retrograde tracer data (***Majka et al., 2020, 2016***). The present resource includes mappings to and from the image spaces used in the aforementioned resources, demonstrated in the present paper by integration with the retrograde tracer data from the MBCA. Further, a diffeomorphic warp between marmoset and human brain has also been added to support qualitative comparison with human brain data. Thus, the BMCA not only provides a new dataset for understanding PFC connectivity, but also a data transfer system for integrating other databases.

This paper describes the post-processing of the data, making it accessible to a broader community of non-imaging experts, and provides access to tools such as the BMCA Explorer and Nora StackApp.

## Results

### The BMCA image processing pipeline

A major part of the work which made the BMCA possible is its image post-processing pipeline, the first for processing serial two-photon tomography images of entire marmoset brains. The pipeline includes fully automated processing of tracer signals, including the detection of the injection site, and the segmentation of anterograde and retrograde tracer signals from the tissue background. It incorporates the mapping of data to a reference image space in high resolution (3 × 3 × 50*μm*^3^) and introduces a new cortical flatmap stack mapping. A flatmap stack is a 3D image representation of the cortex, where the XY-plane defines the position on the cortex surface and the z-direction defines the relative cortical depth. Flatmap stack mappings are extensions of the flatmaps which are part of the MBM atlas and the BMA atlas; Figure 12 shows an example. This section briefly summarizes the data and the pipeline output. Figure 2 shows an overview. Details can be found in the Methods section.

**Figure 2.**
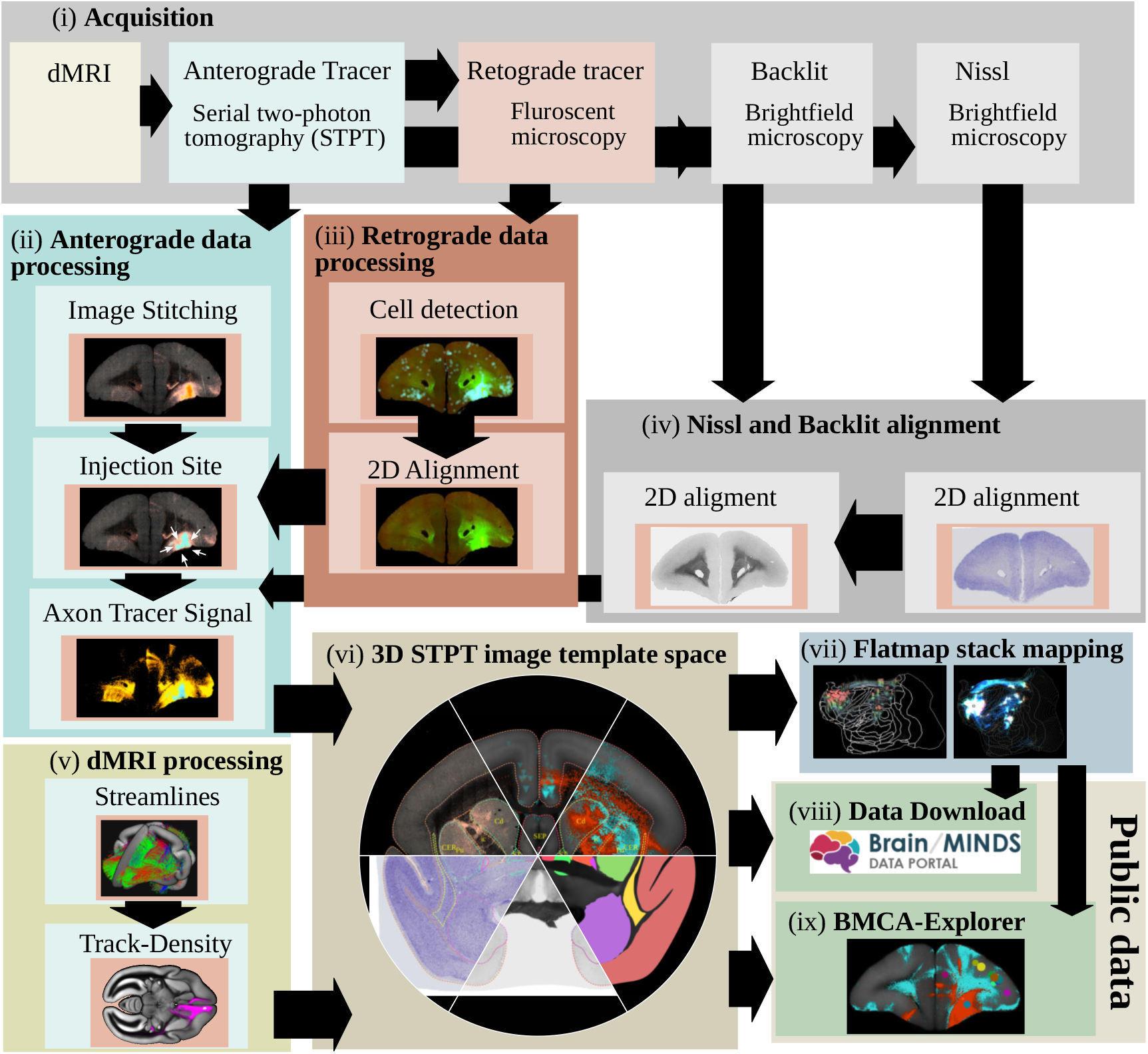
The steps of the BMCA image post-processing pipeline: (i) Image acquisition: After dMRI imaging and automated STPT imaging and sectioning using 2p-tomography, retrograde tracer, Nissl and backlit images are taken. (ii) Processing and analysis of anterograde imaging data, and (iii) retrograde imaging data, respectively. (iv) Automated alignment of Nissl and backlit images. (v) Track-density images are generated from streamlines representing major axon fiber bundles touching the injection site. (vi) All data, including high-resolution microscopy data are mapped to the BMCA 3D brain reference image space. The final steps (vii) are creating the flatmap-stack, (viii) preparing the data for downloading, and (ix) integrating it into the BMCA-Explorer.

#### Pipeline inputs

We obtained data from up to four kinds of image modalities from single marmoset brains: diffusion-weighted MRI (dMRI) for fiber tracking, serial two-photon tomography (STPT) for anterograde tracing, light microscopy for Nissl and backlit images, and fluorescent microscopy for retrograde tracing. The core data of the BMCA are derived from 52 individuals.

In each individual, TET-amplified AAV neural tracer injections (***Hioki et al., 2009***; ***Sadakane et al., 2015***; ***Watakabe et al., 2014***) were placed into a single brain region in the left hemisphere of the marmoset prefrontal cortex; see Figure 16 in the Appendix. Two kinds of ex-vivo full-brain dMRI images were acquired with a 9.4 Tesla MRI animal scanner (Bruker Optik GmbH, Germany). The first image was acquired with the HARDI protocol (b-value of 3000 *s/mm*^2^, isotropic resolution of 0.2 mm, and 128 independent diffusion directions), and the second image as a T2-weighted (T2W) MRI with a 0.1*x*0.1*x*0.2*mm*^3^ resolution. After MRI acquisition, images of 50 µm coronal sections revealing the tracer signal in the entire brain were acquired automatically with TissueCyte STPT (TissueVision, Cambridge, MA). The spatial resolution was 1.385*μm* × 1.339*μm*. We added AAV2retro-EF1-Cre to 19 of the tracer injections, which is a non-fluorescent retrograde neural tracer. After STPT imaging, every 10th section was collected and fluorescently labeled for Cre immunoreactivity. Then, the sections were imaged with an all-in-one microscope (Keyence BZ-X710, Japan). Another set of sections was collected and imaged twice, once before (backlit images, which reveal features of the myelination) and a second time after Nissl staining. We used the same microscope for both backlit and Nissl images (Keyence BZ-X710). The pixel resolution of these sections was 3.774*μm*/px.

#### Pipeline outputs

The purpose of the pipeline was to detect and segment neural tracer signals in the images, perform the fiber tracking, and to integrate all data into a common reference space.

The image processing pipeline automatically performed a 3D image stack reconstruction of the microscopy images, identified the injection site locations, and segmented tracer signals from the background. It then mapped all data, including Nissl and backlit images, to a population average STPT brain image that we used as a template image space. The average STPT brain image was generated by a reiterated registration of 36 subjects (including their left/mirrored versions) using the ANTs image registration toolkit (***Avants et al., 2011***). The pipeline mapped all microscopy images to an isotropic 50*μm* and 100*μm* voxel resolution. In addition, it mapped the data to the STPT template in high resolution. The target resolution for high-resolution data was 3 × 3 × 50*μm*^3^ leading to detailed, co-registered full-brain image stacks with a size of 9666 × 8166 × 800 voxels. The pipeline also automatically integrated measurements such as fiber density and axonal connectivity from the dMRI data. It mapped dMRI data to our template with an isotropic 0.2 mm resolution. It also mapped tracer data to flatmap-stacks, a 3D image representation of the marmoset cortex that extends cortical flatmaps with representations of cortical depth. Finally, it integrated the 3D image stacks into the Nora-Stackapp, and all high-resolution data into the BMCA-Explorer. Both programs are part of the dataset exploration tools that are described in the next section.

### The BMCA-Explorer

The BMCA-Explorer is an online image data viewer that enables visualization of the BMCA data in a high-resolution template space, which is a tremendous advantage for comparative analyses. The viewer shows individual coronal sections of marmoset brain data with an in-plane resolution of 3.0*μm*/px. No previous database viewer could show such high-resolution data in a common reference space.

The Explorer includes anterograde tracer image data obtained in 52 marmosets from the Brain/MINDS project (***Watakabe et al., 2021***). In 19 animals, they are complemented with retrograde tracer data. All data are accompanied by Nissl and backlit sections. For each of the 52 injections, a dMRI tracer density image with directional color encoding is included for qualitative comparison with the neural tracer data. The Explorer also incorporates the data from all 145 retrograde tracer injections from the Marmoset Brain Connectivity Atlas (***Majka et al., 2020***). Further, the BMCA-Explorer provides brain annotations for the Brain/MINDS atlas (***Woodward et al., 2018***), the Marmoset Brain Connectivity Atlas (***Majka et al., 2020***), and the gray and white matter atlases of the Marmoset Brain Mapping project (***Liu et al., 2018, 2020***). It also includes annotations of major cortical and subcortical regions for the current STPT template (***Watakabe et al., 2021***).

Figure 3 shows examples of anterograde tracer data from two different injections. Although the original data are from two different marmosets, we can compare them directly in the same high-resolution space for a detailed comparison. In this example, we can see those axon fibers are highly intermingled in the white matter (white signals), while they are well separated in the striatum or cortex as a whole (3, panel (a)). Interestingly, axon fibers that are completely mixed in the corpus callosum target different cortical regions when entering the cortex (3, panel (c) left). A major advantage of the BMCA Explorer is the axonal-level resolution in the coronal plane. At high resolution, the different trajectories of axon fibers from two samples in the white matter (panel (c) middle) or in the striatum (panel (c) right) can be discerned, which would only be recognized as mixed at low resolution.

**Figure 3.**
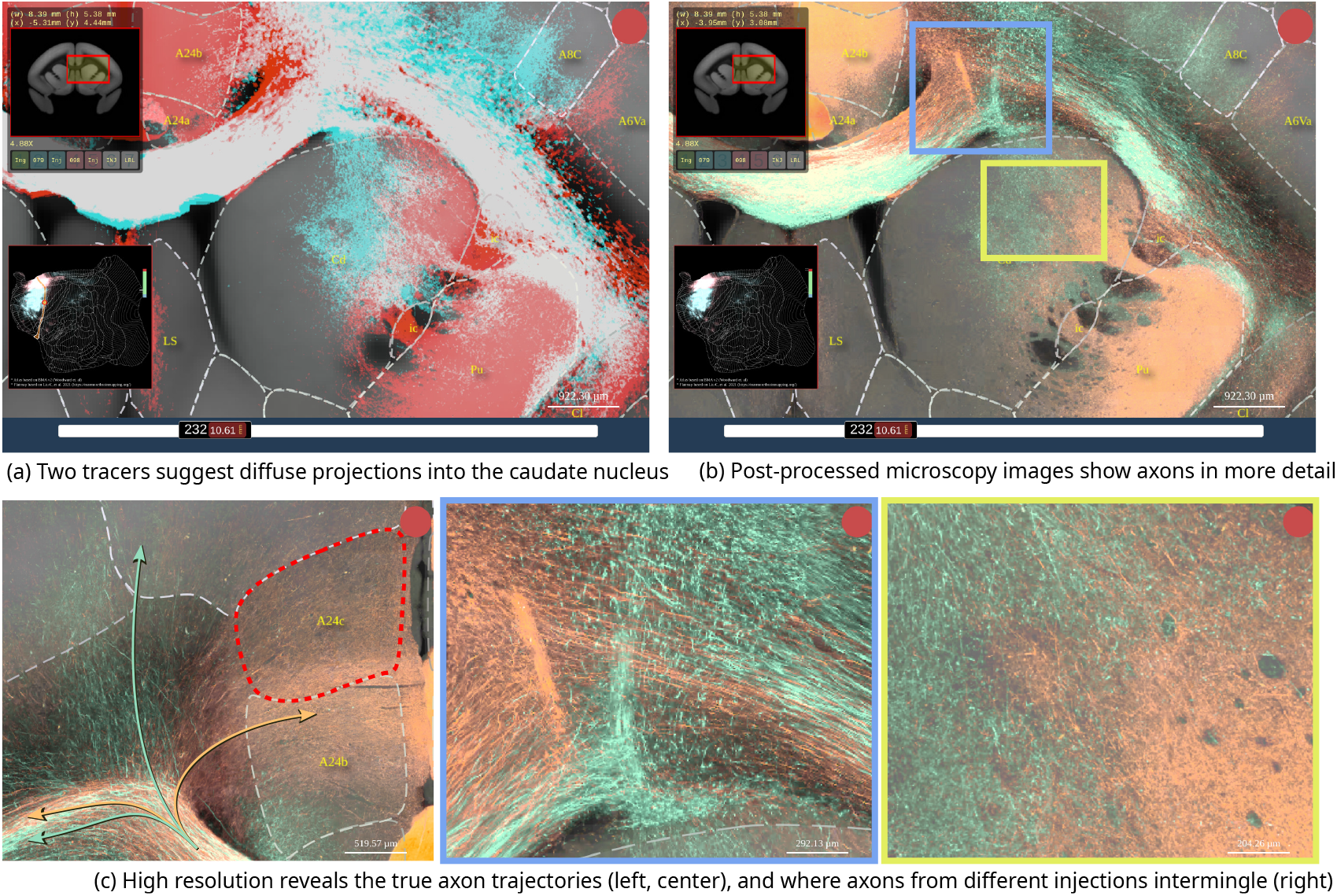
Example screenshots from the BMCA-Explorer demonstrating a virtual overlay of axon fibers originating from two different injections. (a) Tracer segmentation masks for the two samples are shown in red and cyan. In this display mode, the overlap between the two tracers appears in white. Dashed lines are indicating anatomical annotations from an atlas (in this example, the Brain/MINDS atlas). The top left and bottom left panels show the position of the ROI in the coronal section and in the flatmap. (b) Same as in panel (a) except that the figure shows the original image data in two pseudo colors instead of the segmentation mask. (c) High resolution views of panel (b) showing fine details of axonal trajectories.

The BMCA-Explorer is equipped with various tools that facilitate the exploration of the anterograde tracer data. Figure 4 (a) shows two panels on the top left and bottom left for navigation. They show the position of the ROI within the current brain section and its position in the flatmap, respectively. The panel on the right provides access to various atlas annotations, other datasets for comparison, or options to adjust visualization parameters such as contrast and opacity. Available data can be listed and selected by choosing an injection site location from a cortical flatmap, or by selecting a Brain/MINDS marmoset ID; see subfigure (b). In subfigure (c), the synchronization of flatmap overview and the cross-sectional viewer is explained in more detail. Cortical flatmaps are frequently used for visualizing cortical parcellations and connections. However, due to the non-linear deformation and flattening, it is difficult to find corresponding locations in the flatmap and in sections of microscopic image data. The BMCA-Explorer has a flatmap viewer that allows the mapping of flatmap locations to high-resolution microscopy images in real-time, which makes navigating through a flatmap intuitive and fast.

**Figure 4.**
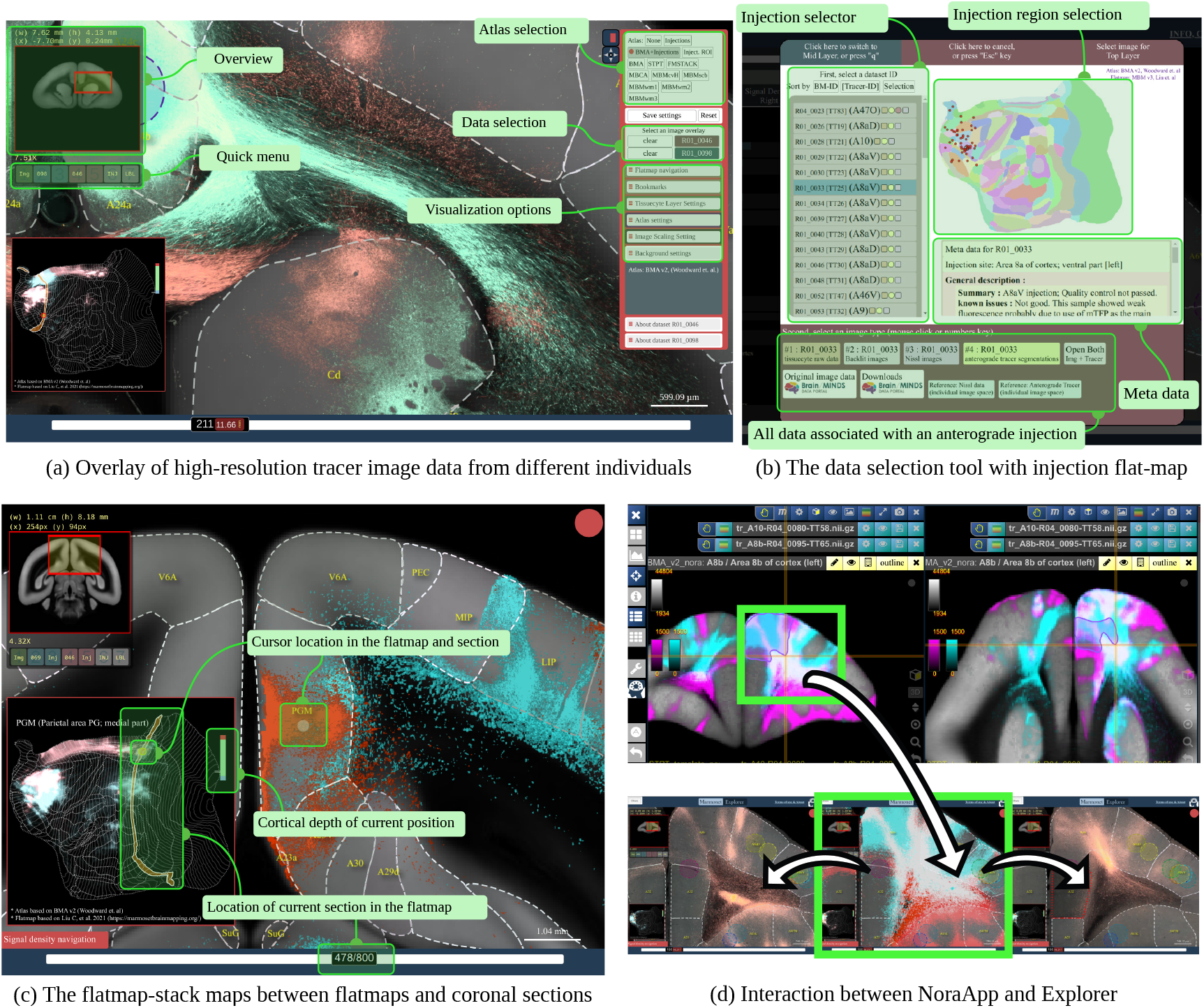
The figure shows the interface of the BMCA-Explorer and The Nora-StackApp. Each example shows two anterograde tracers in the BMCA reference space. (a): The BMCA-Explorer shows high-resolution microscopy images of neural tracers from different individuals in a common image space. Subfigure (b) shows the interface for data selection. (c): The cursor position is shown simultaneously in a cortical flatmap and the current coronal section. (d): The Nora-StackApp viewer can show a number of tracer images simultaneously in 3D which facilitates comparative studies. The viewer supports arbitrary virtual sectioning including sagittal, coronal, or transversal sections, and can interact with the BMCA-Explorer. The same location can be opened in high resolution in the BMCA-Explorer.

#### The Nora-StackApp

Although the original datasets consist of high-resolution images, such images are less suitable for offline use and virtual sectioning. The BMCA provides downscaled isometric volume data for of-fline usage. To support offline exploration, we developed the Nora-StackApp, an image viewer that supplements the BMCA-Explorer with features like virtual sectioning of entire 3D image stacks in a resolution of 100*μm/vox*. The Nora-StackApp is written in JavaScript and is based on the Nora imaging platform (https://www.nora-imaging.com/). The Nora-StackApp facilitates comparative analysis of marmoset brain image data in 3D. For example, once new image data has been warped to the STPT image space, the Nora StackApp can be used to compare the data to all other data in the BMCA. Also, data aligned to any of the three major brain atlases for marmosets can be mapped to the STPT using precomputed warping fields that are part of the BMCA resources. The viewer provides workspaces based on the BMA, BMCA and the MBM atlases and can overlay numbers of tracer images and fiber density maps simultaneously. Figure 4 (d) shows a screenshot. The 3D image stacks provide a global picture of the neural architecture in low resolution. At any time, details can be inspected by opening a coronal section in high resolution at the exact same position in the BMCA-Explorer.

### Comparing anterograde neural tracer with dMRI tractography using BMCA data

Diffusion MRI is widely used for studying primate brain connectivity in vivo. It is thought to reflect the anisotropy of axonal fiber structures. However, the estimates are imperfect (***Thomas et al., 2014***; ***Girard et al., 2020***). The BMCA provides anterograde tracer data showing axonal projections from the injection site. When combined with dMRI, it can be used as a “ground truth” for comparison with dMRI-based tractography.

For instance, areas A32 and A8aV are two distinct regions in the PFC where the tracer signals show non-overlapping pathways. Interestingly, a comparison of the two pathways showed that they are also separated in the corpus callosum and the internal capsule. The spatial gap between the two injection sites, and the non-overlapping pathways in close proximity, make these two regions a perfect example for comparison with dMRI fiber tractography.

Figure 5 shows the tracer signal in comparison with the fiber density maps generated from dMRI fiber tractography. Since we have multiple injections into both regions, we have merged the tracer signal of the injections in each injection site by taking the maximum of the normalized tracer signals. Further details regarding the tracing and the fiber tractography method are explained in the Methods section.

**Figure 5.**
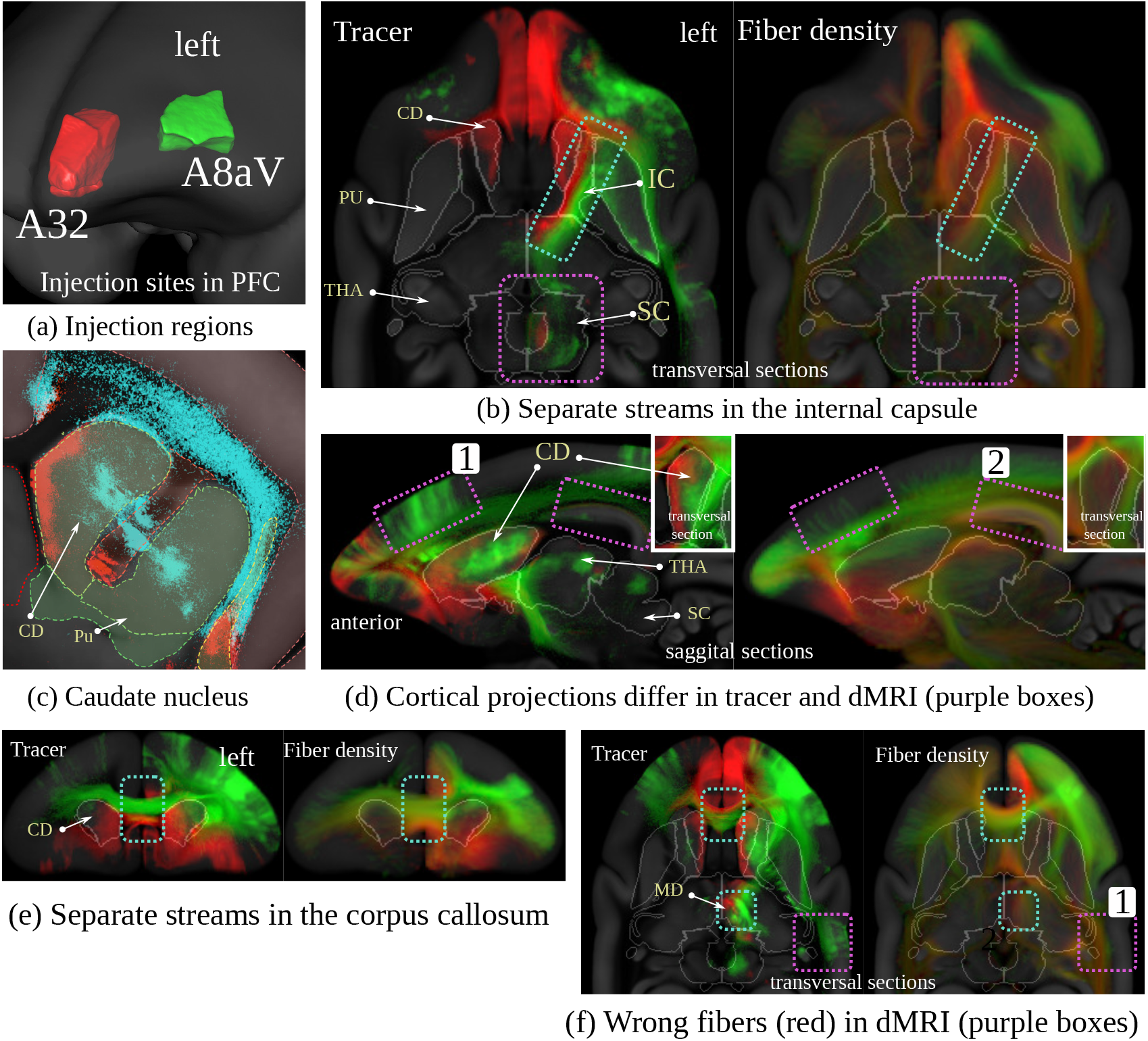
Visual comparison between dMRI-based fiber density and anterograde neural tracers originating in two distinct regions in the marmoset PFC. Each of the two colors represents the maximum over all anterograde tracers that have been injected into one of the two regions in the marmoset PFC (A32: red, A8aV: green (cyan in (c))). The similarity between the images suggests that dMRI reflects real brain connectivity (cyan boxes with dashed borders in (b),(d),(e) and (f)) but also shows evidence of the relative imprecision of dMRI data in terms of specificity and sensitivity (violet boxes with dashed border).

There are remarkable similarities between the dMRI and tracer data. Figure 5 (b) shows that both tracer and fiber tracts pass the internal capsule in segregate streams. The tracts have strong projections into the mediodorsal thalamic nucleus (MD). There has been strong evidence from dMRI tractography that A32 projects anteromedially while A8aV projects posterolaterally (***Phillips et al., 2019***), which could be confirmed in our comparison of tracer and dMRI tractography (Fig. 5 (f) and Figure 15 in the appendix). Tracts from both regions also pass the corpus callosum in separate streams (Fig. 5 (e) and (f)). The tract from A8aV runs ventroposteriorly in the corpus callosum, while the tract originating in A32 ventralposteriorly.

However, there are also clear discrepancies between the two sets of results. Figure 5 (c) shows diffuse tracer projections into the caudate nucleus (CD). The image shows a close-up view of the segmented tracer of two examples, one for each injection area. A32 projects medially, and A8aV laterally. These projections are not present in the dMRI tractogram. Similarly, we can observe thin, diffused connections from the thalamus projecting strongly into the superior colliculus (Figure 5 (b)), which are also absent in the dMRI data. It is also occasionally difficult to reproduce cortical projections correctly. For example, the region highlighted by box 1 in Figure 5 (d) shows strong cortical projections in the tracer signal, which are underrepresented in the data derived from dMRI. Conversely, box 2 shows strong cortical projections in dMRI that are not present in the tracer data. A similar observation can be made in box 1 in Figure 5 (f), where dMRI tractography suggests fiber bundles which are not supported by the tracer data.

These direct comparisons add to the evidence of the relative imprecision of dMRI data in terms of specificity and sensitivity, which has been proposed based on comparison of tracer and tractography data in the macaque brain (***Thomas et al., 2014***).

### Integrating retrograde tracer data of the Marmoset Brain Connectivity Atlas

The Marmoset Brain Connectivity Atlas (MBCA, https://www.marmosetbrain.org/) provides postprocessed image data for 145 retrograde tracer injections. Retrograde tracers can reveal back-projection; making them a valuable counterpart to our BMCA anterograde tracer data, given the general reciprocity of corticocortical connectivity (***Rockland, 2015***). The data contain the location of cell bodies in the Paxinos stereotaxic reference space (***Paxinos et al., 2012***). We mapped all 145 data sets to our BMCA template image space. The matrix in Figure 6 (b) shows the normalized cross-correlation of the anterograde tracer signal and cell density in the cortex between pairs of flatmap-stack data. Data were paired with respect to the closest injection site distance with respect to the STPT template space. Subfigure (a) shows the similarity of the anterograde tracer data as a reference.

**Figure 6.**
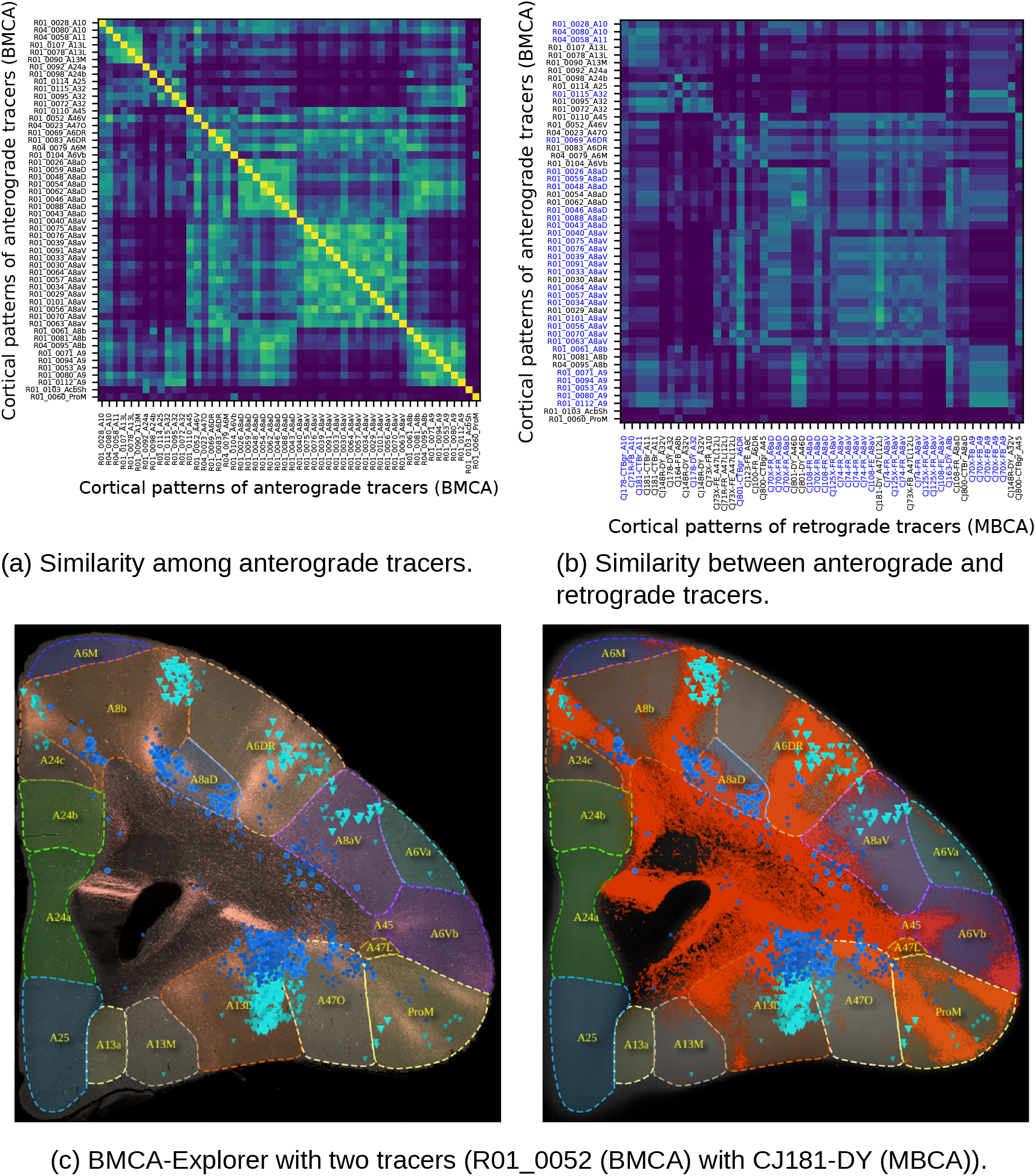
We integrated retrograde tracer data from the Marmoset Brain Connectivity Atlas (MBCA) into the BMCA. The top matrices show the visual similarity of tracer data in the cortex of marmosets (based on normalized cross-correlation). Subfigure (a): similarity between all pairs of anterograde tracers as a reference, and for (b), pairs were formed between BMCA anterograde tracers and MBCA retrograde tracers based on the distance of the nearest injection site. Subfigure (c) shows an example of notable overlap between a pair of data from both projects with similar injection sites. The left image shows the retrograde tracer (cyan: supragranular, blue: infragranular) over the anterograde tracer signal. The right image shows the retrograde signal over the segmentation of the anterograde tracer (red).

The data were integrated into the BMCA-Explorer and the Nora-StackApp. We also mapped cell density onto flatmap stacks. The data suggest that anterograde tracers and retrograde tracers show similar projection patterns, but reveal the important differences in their laminar patterns, which is essential for the definition of feedforward and feedback connection patterns (***Theodoni et al., 2022***; ***Markov et al., 2014b***). The similarity between the tracers suggests that the MBCA retrograde data are well aligned with our STPT template image space. Sub-figure (c) shows an example in the BMCA-Explorer for two nearby injections in the prefrontal cortex. It illustrates an example of remarkable spatial correspondence between the two tracer patterns.

### Mapping human anatomy to marmoset

One overarching goal is to incorporate knowledge gained from marmoset brain anatomy and structural connectivity into our understanding of the human brain. However, it is illusive to believe that there is a one-to-one mapping from human to marmoset brain, which reflects all structural and functional aspects (***Sneve et al., 2019***). But, unquestionably, there are strong similarities. Towards this goal, we present here a diffeomorphic warp from human to marmoset anatomy and vice-versa, which will enable scientists to relate their marmoset findings to human anatomy.

When looking at marmoset and human brain anatomy, in particular, the white matter morphology is similar among the two species. Hence, we aimed for co-registering human diffusion MRI (living in MNI2009b atlas space) to marmoset HARDI average data residing in our marmoset template space. Registration is solely based on the anatomical/structural features regardless of functional aspects. Details of the registration approach can be found in the method section. In Figure 7 we show qualitative comparisons of the obtained registration. The correspondences (based on colored fractional anisotropy) in white matter match astonishing well (Figure 7 (a). The T2-weighted MRI images in Figure 7 (b) show that details are not perfectly matching, however, the gross anatomy is congruent. Figure 7 (d) shows an overlay of the average of all anterograde tracer segmentations which were warped to human MNI space. One can see that the white matter structures (like the anterior capsule) are well-matched. Also, the midbrain pathway passing the subthalamic nucleus is in place. Figure 7 (c) shows the overlap from 14 regions (quantified by the DICE coefficient) of the human JHU (JJHU-ICBM-labels-1mm.nii.gz) (***Hua et al., 2008***; ***Wakana et al., 2007***; ***Mori et al., 2005***) white matter atlas with the marmoset white matter pathways atlas of the MBM project (***Liu et al., 2020***). While there are still some problems (like the Tapetum and the medial lemniscus) the overall agreement is promising.

**Figure 7.**
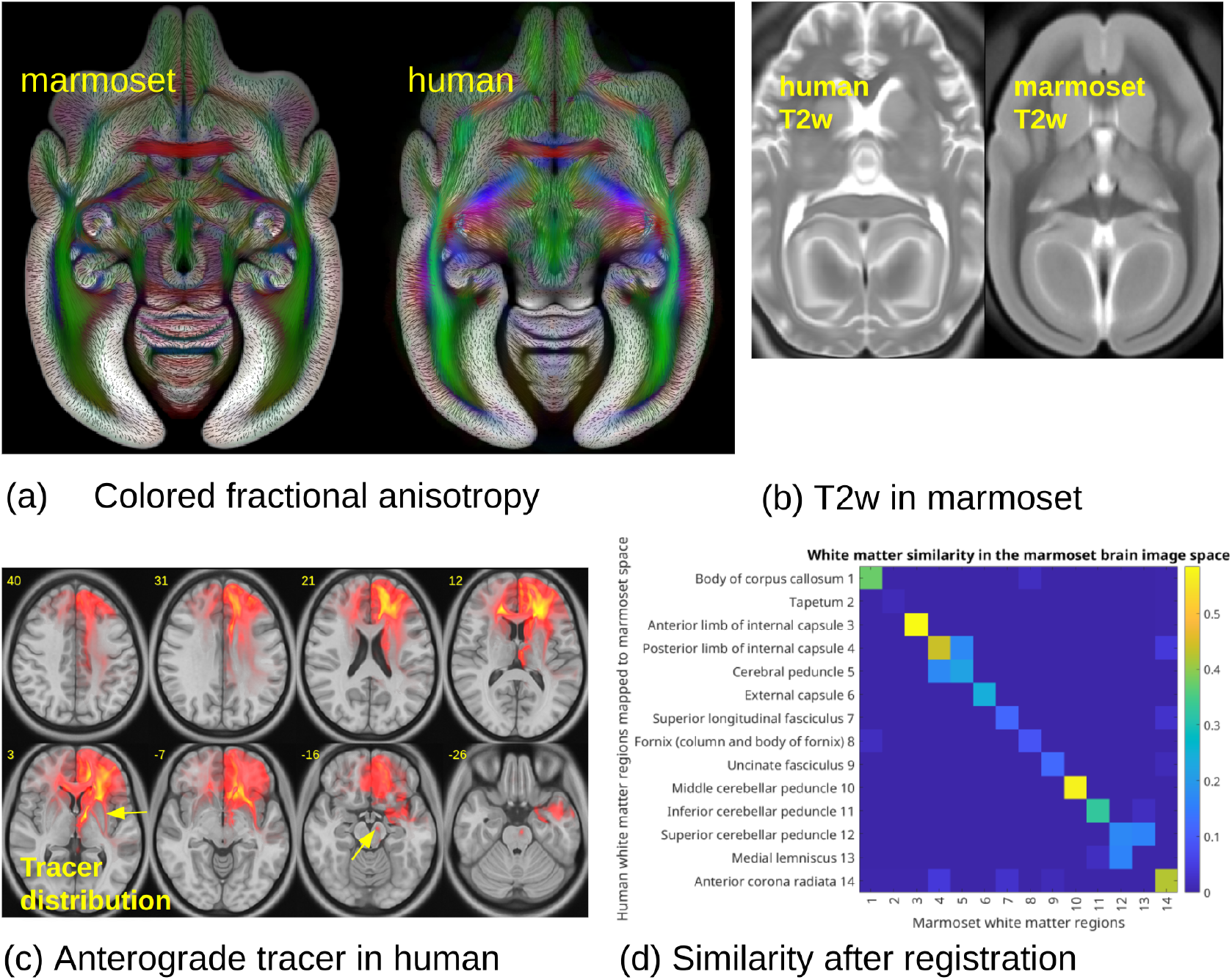
Sub-figure (a) shows fractional anisotropy in human and marmoset brain dMRI images in the BMCA template space. Figure (b) shows images of a human brain and a marmoset brain in the BMCA template space. Figure (c) shows a tracer density mapped to the human MNI image space, and (d) shows the DICE similarity of major white matter regions between a marmoset and the human brain white matter atlas after mapping the latter from human to the marmoset brain.

## Discussion

The work in this paper is part of Japan’s Brain/MINDS project (***Okano et al., 2015***; ***Okano and Mitra, 2015***; ***Okano et al., 2016***). The project is working on the construction of an integrated, multiscale structural map of the marmoset brain from data acquired using several imaging modalities such as 2photon imaging, in situ hybridization (***Kita et al., 2021***), and dMRI. The BMCA tools described here allow exploration of the first publicly available multimodal dataset of anterograde tracer injections in a primate brain. The integration of neuroanatomical tracers with structural MRI allows the user to navigate bidirectionally between macroscopic anatomical information, obtained by MRI, and cellular-level neuroanatomical information obtained by tracers and histological techniques. In addition, the BMCA allows direct comparisons between anterograde and retrograde tracer injection data, due to the integration of datasets from the Marmoset Brain Connectivity Atlas (***Majka et al., 2021***). Whereas the emphasis here is on the connectivity of marmoset PFC, current work is aiming to extend the data with anterograde tracer injections covering other regions of the cortex. Other planned features include open access raw dMRI image data from each individual animal, and data from disease model marmosets.

### Relation to previous work

The development of the BMCA is part of the international trend towards open-access resources for the exploration of brain connectivity. Integration of connectivity datasets into multimodal platforms has recently been identified as a priority area for the advancement of translational neuroscience (***Milham et al., 2022***), and the present resource addresses this need. In this regard, the BMCA extends and complements capabilities offered by other online resources. For example, the Human Connectome Project compiles an extensive amount of such structural and functional neural data of the human brain (***Marcus et al., 2011***). However, the acquisition of large-scale structural connectivity data is limited to dMRI imaging techniques. For animal models, tracer techniques are frequently used to map neural connectivity in more detail (***Zingg et al., 2014***; ***Markov et al., 2014a***), with the Allen Mouse Brain Connectivity Atlas (***Oh et al., 2014***; ***Kuan et al., 2015***) and the Marmoset Brain Connectivity Atlas (***Majka et al., 2016, 2020***) offering two examples where the results of a large number of tracer injections is made publicly available, and accompanied by an average template brain, brain annotations, and tools for visualization and exploration. The Allen Mouse Brain Connectivity Atlas provides anterograde tracer data in the mouse brain, which have been acquired with an STPT system (similar to the present BMCA). In contrast, the Marmoset Brain Connectivity Atlas reconstructs data from cortical retrograde tracer injections from histological sections of the marmoset brain, followed by 3-dimensional reconstruction and registration to stereotaxic reference space.

Overall, there still exist only a few integrative tracer databases, to some extent due to the fact that the systematic mapping, processing, and visualization of imaging data are labor-intensive and costly. Alternative approaches, such as the CoCoMac project (***Bakker et al., 2012***; ***Stephan et al., 2001***), aim at accumulating and integrating the output of various research studies to get a better picture of global brain connectivity. However, this relies on heterogeneous data sources and lacks access to ready-to-use image data.

### Originality and significance

The prefrontal cortex plays a key role in the more complex aspects of primate cognition, and its dysfunction is involved in various psychiatric disorders. In this setting, the common marmoset monkey is considered to have great potential as a model organism for disorders of cognition (***Okano, 2021***; ***Miller et al., 2016***; ***Belmonte et al., 2015***). However, publicly available post-processed image data of anterograde axonal tracer injections covering the marmoset PFC have not been available.

The STPT technique enables accurate 3D tomographic reconstructions of the acquired image stacks and tracer patterns. This allows the patterns to be registered into a common template space, which is a prerequisite for any sound comparison and quantitative evaluation. The BMCA tools give access to this rich repository of high-quality data, and support its integration to other Marmoset brain projects such as the Marmoset Brain Connectivity Atlas (***Majka et al., 2020***) and the Marmoset Brain Mapping project (***Liu et al., 2021***). It is also the first time that a diffeomorphic warp between marmoset and the human brain is made publicly available, an essential step towards interspecies data translation.

### Integration of different modalities of structural connectivity

Diffusion MRI is currently the most widely used technique for studying brain connectivity. dMRI provides a key link to neuropsychological and neurosurgical practice, in particular, due to its in vivo applicability. The main image features of dMRI are based on relatively large and oriented axonal fiber bundles, which create an anisotropy of the dMRI signal. However, estimates of true structural connectivity based on this technique are imperfect (***Thomas et al., 2014***; ***Girard et al., 2020***), and depend to a large extent on postprocessing steps (***Maier-Hein et al., 2017***). Anterograde injection studies are important, because they can be used to validate connectivity measures based on diffusion MRI (HARDI) (***Jiang et al., 2006***; ***Tournier et al., 2012***). The BMCA integrates a HARDI population image to foster the comparison between tracer and structural dMRI (***Goulas et al., 2019***; ***Girard et al., 2020***; ***Gutierrez et al., 2020***). In this context, the BMCA provides a rich platform to enable future studies aimed at refining and validating dMRI, by providing simultaneous “ground truth” datasets, and access to histological information. Figure 5 shows that the similarities in topology between tracer injection patterns and dMRI fiber densities can be remarkable. Although the bundles of different injections run fairly close to each other, the dMRI fiber densities stay consistently apart, which suggests that assumptions about topological preservation, which most tractographic approaches rely on, are generally valid. However, cellular tracer data allow estimates of the directionality of the connection, which is not recoverable from dMRI data.

Further, the BMCA integrates anterograde tracer data with retrograde tracer data. Our recent study has shown that signals from combined anterograde and retrograde tracer injections correlate well in the PFC, suggesting a strong correlation between projection patterns and back projection patterns (***Watakabe et al., 2021***). However, there are consistent reports of non-reciprocal pathways in both the macaque and marmoset brain (***Theodoni et al., 2022***), and the combination of both tracer modalities is the best approach to further investigate this issue. In addition, the laminar patterns of both cell bodies and terminals are critical for establishing patterns of hierarchical flow of information in multi-areal pathways (***Felleman and Van Essen, 1991***). To encourage further investigation of such relationships, we supplemented our data with the large set of retrograde data from the Marmoset Brain Connectivity Atlas. Our preliminary comparison with the MBCA data supports our observation of spatially correlated tracer patterns and reveals different laminar patterns for cell bodies and terminals within cortical columns.

### Importance for translation of knowledge to human application

The importance of non-human primates for the translation of biological findings to human health applications has been repeatedly emphasized (***Wang et al., 2020***; ***Milham et al., 2022***). Indeed, primate models like the marmoset have gained popularity because of their similarity in brain structure with the human brain, and cognitive abilities, which allow for informed analysis of various brainrelated diseases such as autism, schizophrenia, or dementia. However, to directly relate the image information from different species using computational mapping tools, such as between the marmoset and the human brain, remains a challenge. The present study proposes a direct mapping between imaging data of the marmoset and human brain, based on matching similar structures in the white matter. Although preliminary results have shown similarities that suggest the value of this approach, it is important to take into account differences in the structure of the gray matter, such as the larger number of subdivisions in the human cortex (***Glasser et al., 2016***). Further efforts are needed to improve and validate the matching in terms of structure and function, perhaps taking into account the known patterns of differential expansion as a function of brain volume (***Chaplin et al., 2013***) and the likely effects of this on patterns of connectivity (***Rosa et al., 2019***), will be a rich area of future study.

## Materials and methods

### The BMCA image data-set

This section briefly describes image acquisition. Details regarding the injections, the neural tracer, and the acquisition can be found in the methods of the previous report (***Watakabe et al., 2021***). The data set comprises the multimodal image data of 52 individuals with 52 anterograde tracer injections. The injections were placed into 21 disjunctive brain regions in the left hemisphere of the marmoset prefrontal cortex. Figure 16 in the appendix shows the location of the 52 injections. The acquisition took place in five steps. The TET-amplified AAV tracer system contained a mixture of clover and 1/4 amount of presynapse targeting mTFP1, a fluorescent anterograde neural tracer. For 19 brains, the tracer was injected in a mixture with AAV2retro-EF1-Cre, a retrograde tracer.

First, postmortem, an ex-vivo full-brain diffusion-weighted MRI (dMRI) was imaged with a 9.4 Tesla MRI animal scanner (Bruker Optik GmbH, Germany). The dMRI images were based on the HARDI protocol with a b-value of 3000 *s/mm*^2^ with an isotropic image resolution of 0.2 mm and 128 independent diffusion directions. In addition, T2-weighted (T2W) images were taken with a spatial resolution was 100 um × 100 um × 200 um.

Next, the entire brain with the fluorescent anterograde tracer signal was imaged by fully automated Tissuecyte 1000 and Tissuecyte 1100 serial two-photon tomography (TissueVision, Cambridge, MA). For STPT, the entire brain was embedded into agarose and mounted under the microscope. After imaging the block face, automated vibratome sectioning was performed. These steps were repeated automatically until the entire brain was imaged. The coronal in-plane image resolution of the raw data was 1.385*μm* × 1.339*μm*^2^ with in total of about 19000 × 16000 pixels. The image of each section contained three channels: The first channel best represents the auto-fluorescent background (the entire brain structure) and the second channel is the tracer signal. There was a large difference in tracer intensity within and outside the injection site. To capture weak tracer signals, the dynamic range was sacrificed in the injection site such that the signal in the injection site was saturated in the first two channels. For compensation, the third channel was used to represent the details (the infected cell bodies) in the injection site. Figure 9 (a) shows an example.

Every 10th section, which corresponds to a 500 *μm* offset, was recovered and fluorescently immunostained for Cre. Fluorescent images were captured with an all-in-one microscope (Keyence BZ-X710). The in-plane resolution was 3.774*μm*/px. The images contain two color channels. The first channel contains the signal of the retrograde tracer signal (cell bodies). In the second channel, the anterograde tracer signal was also captured by the Tissuecyte microscope. In addition, the second set of slices, also in a 10-section interval, was collected. The sections were imaged twice before (backlit) and after Nissl staining with the same microscope (Keyence BZ-X710) with a pixel resolution of 3.774*μm*/px.

### The processing pipeline

This section describes details regarding all post-processing methods. Figure 2 outlines the processing pipeline. The image post-processing part of the pipeline works in a fully automated manner and does not require manual interaction. The pipeline was written in a mixture of python 3 (www.python.org) and Matlab (MathWorks) code. Each pipeline step was executed by a dedicated script running on an Ubuntu Linux cluster system. The entire pipeline was orchestrated by a pipeline database system written in python that was keeping track of data dependencies and was able to lunch scripts in parallel using the SLURM workload manager (www.schedmd.com).

All image data has been aligned to our BMCA template space, a left-right symmetric population average template of a marmoset brain with an isotropic resolution of 50*μm*. For image alignment, we used the ANTs image registration. If not stated otherwise, we used a multi-scale affine image registration followed by a multi-scale deformable SyN registration (***Avants et al., 2011***) with normalized mutual information as a metric.

#### STPT image stitching

The image stitching was done in Matlab based on in-house code. The TissueCyte microscope generates a large number of image tiles. The size of each image tile has been set to 720 × 720 pixels with a spatial in-plane resolution of 1.385*μm* × 1.339*μm*^2^. The tiles were provided as 16bit tiff files. The microscope outputs the offset for each image tile in plane-text as 3D world coordinates with micrometer resolution. The coordinates are sufficiently precise to allow the reconstruction of an entire image section by aligning and fusing all image tiles according to their world coordinates. We set a small overlap between adjacent image tiles (about 80 pixels) and cropped 50 pixels from the image tile boundaries before fusing them using a linear blending function. The size of an entire brain section was approx. 19000 × 16000 pixels.

Distortions in the microscope’s optical path created an inhomogeneous vignetting effect in the tile images. Hence, before stitching, we applied an intensity correction. We estimated the shading field by averaging over a large set of image tile samples and divided each tile by the result. Intensity correction for tile images was only applied to the first two channels (background and tracer). The third channel, which has a clear signal around the injection site but a low contrast and a bad signal-to-noise ratio elsewhere was excluded from the correction due to the small number of tile samples with meaningful content. Further details regarding the correction algorithm can be found in our technical report (***Skibbe et al., 2019***). An example before and after stitching is shown in Figure 9 (b).

For all three image channels, we created 3D image stacks with an isotropic image resolution of 50*μm*. The image stacks were saved in the NIfTI file format (https://nifti.nimh.nih.gov/). The full resolution image sections were stored as 16bit lossless PNG images.

#### Injection site location

The pipeline locates the injection site in two steps. Figure 9 (c) shows an example. First, we located its rough position as the brightest connected structure in the 3D image stack of the third STPT color channel. In that channel, the cells in the injection site appear bright, while there is almost no signal outside the injection site. For the localization, we used Matlab. We applied Gaussian smoothing, followed by the application of an intensity threshold (half the maximum intensity in the image) and connected component analysis. We determined the volume of the injection site as the largest connected component.

In a second step, an artificial neural network screened all full-resolution 2D STPT brain sections for infected cell bodies. To speed up the process, the screening only took place for that part of the 2D image sections that intersected with the volume of the injection site, which was determined in the first step. The network architecture was a 2D U-Net, a convolutional neural network for biomedical image processing (***Ronneberger et al., 2015***). We trained the network on images with 512*x*512 pixels to map STPT images to probability maps for the locations of the centers of cell bodies. Local maxima in the probability maps that exceeded a probability of 0.5 were considered cell locations.

We created a training data set with images of 6068 manually annotated cell bodies that appeared in 44 different 2D images. We selected the images from 10 different marmoset brains. The U-Net was based on the original implementation, with a depth of four. Most upper layers had 64 features after the first convolution layer. The number doubled after each pooling operation to up to 512 features. We further used drop-out and batch normalization (***Ioffe and Szegedy, 2015***; ***Srivastava et al., 2014***). The training procedure augmented the image with deformations and changes in intensity and contrast. Details regarding the architecture and a performance evaluation can be found in our technical report (***Skibbe et al., 2019***).

#### Anterograde tracer segmentation

This step takes the raw STPT image sections as input and segments the anterograde tracer signal from the background. This was done by applying a 2D U-Net to the data. The network took image patches combining the first two STPT image channels as input and returned a tracer signal probability map as output. Both input channels show the auto-fluorescent background, but the neural tracer signal was significantly brighter in the second channel. The difference helped the network to better distinguish the tracer-positive pixels from the background. The pipeline applied the network to all image sections. Figure 9 (d) shows an example.

The pipeline generated two kinds of image data from the segmented data. The anterograde tracer density, and the normalized anterograde tracer signal intensity. The tracer *density* is a 3D image stack in which voxels represent the amount of tracer positive pixels in an 50*x*50*μm*^2^ area in a raw 2D image section. The normalized tracer signal *intensity* takes the actual tracer intensity into account. The neural tracer is much brighter in the second channel than in the first channel, while the background appears similarly bright in both channels. We obtained the signal intensity by subtracting the first channel from the second channel. We then normalized the intensity with respect to its strongest signal outside the injection site. The injection site itself was excluded from the calculation because of the saturated signal in the injection site, and thus values within the injection volume did not represent a meaningful quantity for normalization. Figure 8 shows an example of 3D reconstruction of tracer density.

**Figure 8.**
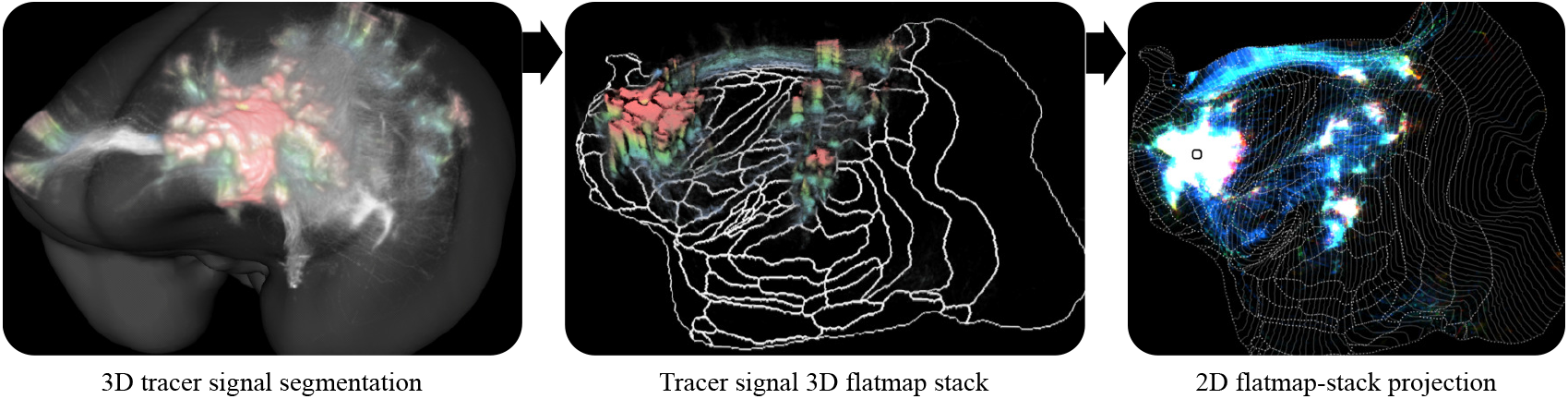
A 3D reconstruction of an anterograde tracer density. The intensities in the cortex have been colored according to cortical depth. From left to right: The signal in the STPT template image space, the signal in the left hemisphere mapped to a 3D flatmap stack, and the 2D projection of the flatmap stack.

**Figure 9.**
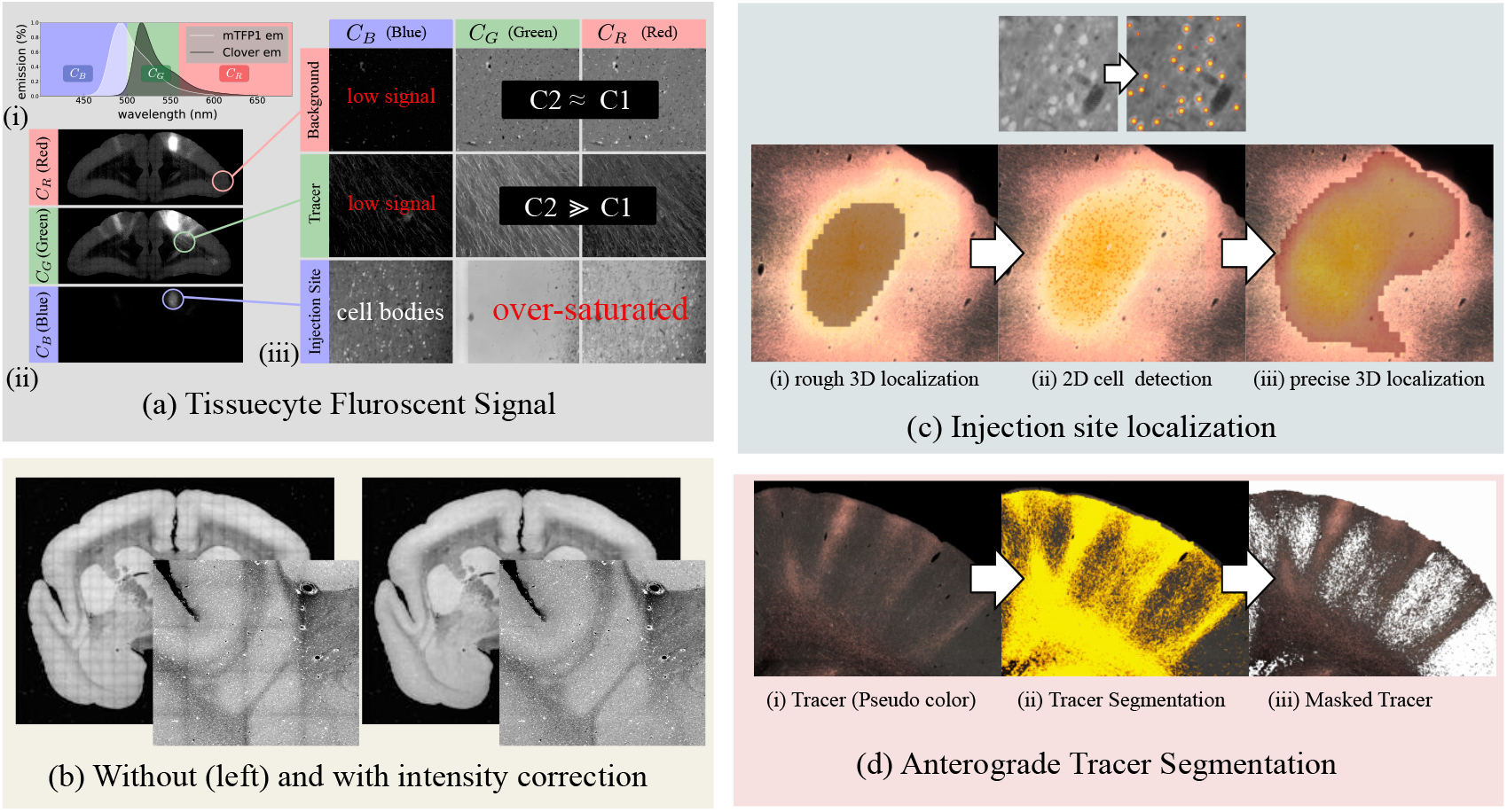
(a) The fluorescent emission profile of the anterograde tracer. (b) A coronal section of the STPT microscope (first channel) before and after intensity correction. (c) Illustration of the injections site localization, and (d) the anterograde tracer segmentation.

The data for training the U-Net were generated in a semi-supervised way. We applied a threshold to the tracer intensity to generate a large set of labeled brain image sections. We manually screened and selected about 600 image sections from 20 different marmoset brain image stacks for training. Various structures with bright signals that were not part of neurons, like blood vessels, were manually annotated as “explicit negative” examples. We added an extra penalty to the detection of such structures during training. We generated a training set of 12000 smaller image tiles from the labeled data. We used the same kinds of data augmentations as for the cell body location. Further details can be found in (***Skibbe et al., 2019***).

#### Retrograde tracer segmentations

Figure 10 illustrates the acquisition and post-processing of retrograde tracer signals. All image sections of the retrograde tracer had two color channels, where the first channel (red) contained the cell bodies of the retrograde tracer. The second channel (green) contained the anterograde tracer signal that is also visible in the Tissuecyte microscope (STPT). We utilized the first channel to localize the cell bodies of retrogradely infected neurons and exploited the second channel to align the image to its corresponding Tissuecyte section.

**Figure 10.**
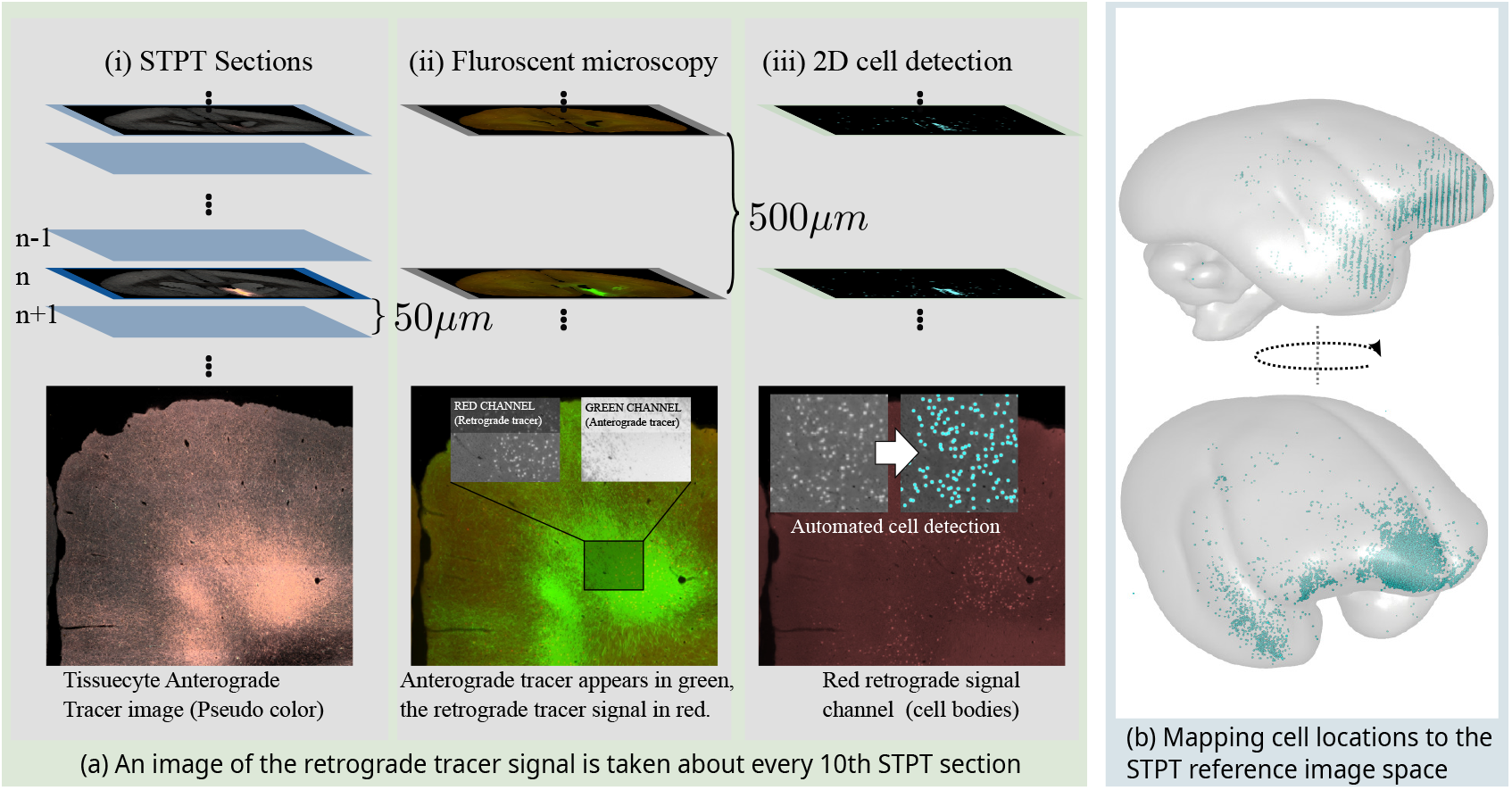
Retrograde tracer segmentations. (a) Every 10th Tissuecyte section, which corresponds to a 500 *μm* offset, was recovered and fluorescently immunostained for Cre. Fluorescent images were captured, and a convolutional neural network was applied to detect cell bodies in the image. (b) The cell body locations were mapped to the BMCA template image.

Similar to the detection of cell bodies in the injections site for anterograde tracers, a U-Net was trained and used for cell body detection. The network took patches (sized 512 × 512) as input and was applied to all image sections. Local maxima in the results with a probability larger than 0.5 were considered as detection.

The pipeline registered the second channel with the anterograde tracer signal to the corresponding Tissuecyte section using ANTs. The same image transformation was applied to the first channel and the location of detected cell bodies.

For training the U-Net, about 20,000 patches were randomly sampled from 380 manually annotated image sections (roughly about 20,000 training patches).

#### The BMCA STPT template space

For data integration and investigation, all imaging data was automatically normalized to a volumetric STPT average template with left/right hemispherical symmetry. The auto-fluorescent back-ground signals (the first channel) of individual STPT images were used for registration and for computing the population average image. Figure 11 (a) shows the template. While averaging, values were inversely weighted by their tracer intensity so that image data that was dominated by signals of neural tracers was suppressed. Areas with missing tissue were excluded as well. The template was generated by a reiterated registration of 36 subjects (including their left/mirrored versions); see Figure 11 (b). The STPT template has an isotropic resolution of 50*μm*.

**Figure 11.**
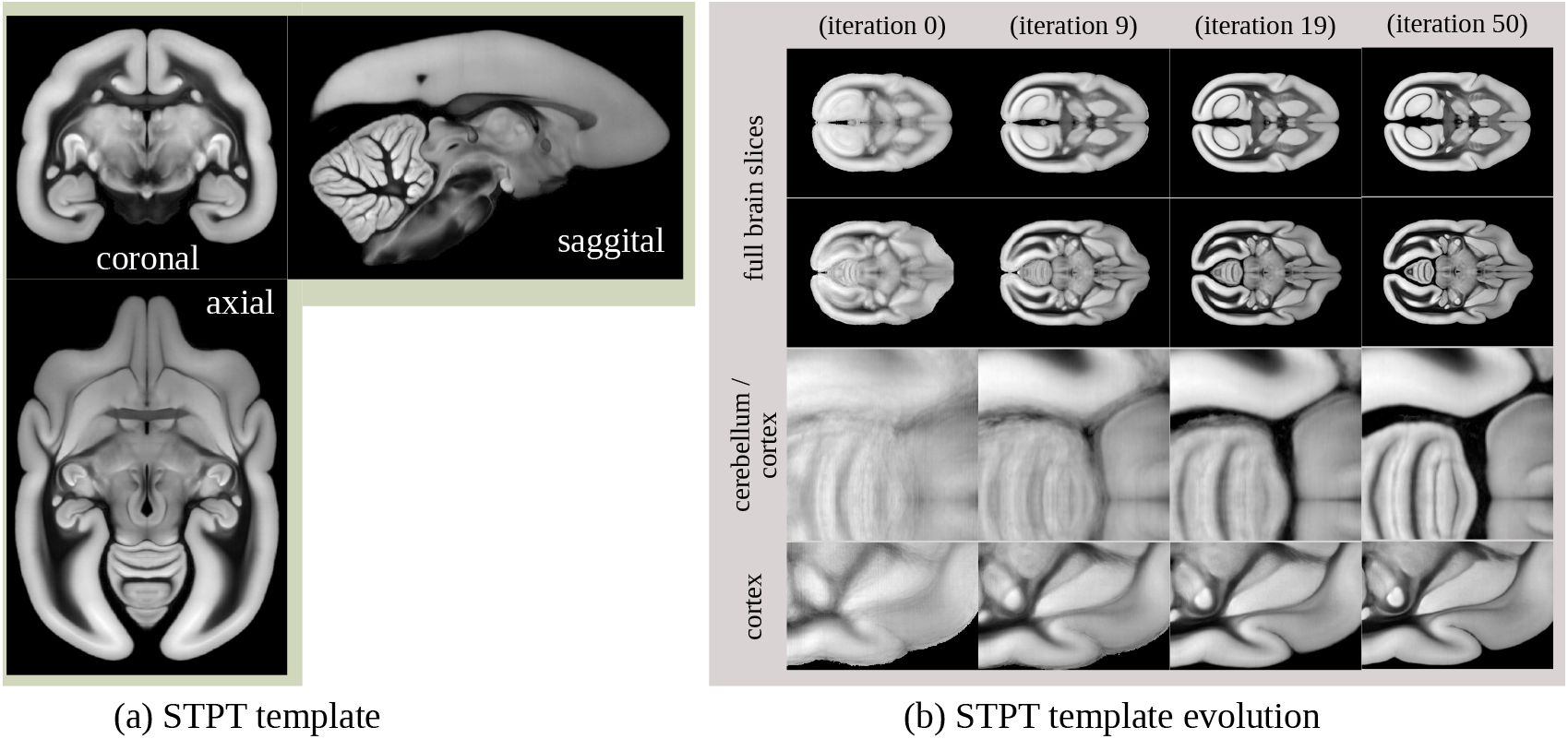
The BMCA Common Coordinate Framework is defined by a population average STPT template image. The STPT template was generated by reiterating the registration of all subjects.

**Figure 12.**
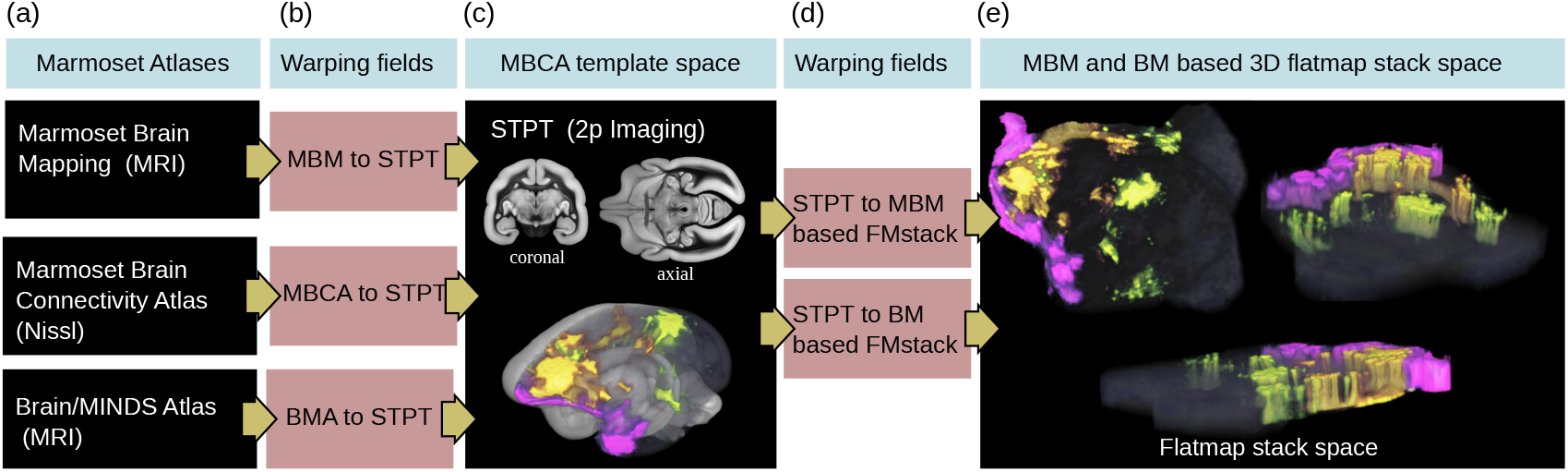
The BMCA provides the files for mapping between all major marmoset brain coordinate frameworks. It further can map cortical 3D image data to a flatmap stack using the publicly available ANTs image registration toolkit. This example shows the combined mapping of three anterograde tracer images from a 3D brain image to a flatmap stack.

The spatial resolution of our STPT image sections was sufficiently high to map the microscopy image sections to our template in high resolution. For web deployment, microscopy images were mapped to our template with a remarkably high target resolution of 3.0 × 3.0 × 50*μm*^3^. The resulting image stacks contained 9666 × 8166 × 800 voxels. All images have been processed and compressed to make them suitable for fast web exploration using either PNG, JPG, or the modern AVIF image file format.

The STPT template was accompanied by a label image annotating the cortex and major subcortical structures such as the thalamus, caudate nucleus, internal capsule, putamen, or hippocampus.

#### Atlas mapping

We computed the transformation fields to map between the MBCA reference space and three major marmoset brain atlases which are, the Marmoset Brain Mapping atlas (MBM) version 2 and 3 (***Liu et al., 2021, 2020***), the Marmoset Brain Connectivity Atlas (MBCA) (***Majka et al., 2016, 2020***), and the Brain Minds Atlas (BMA) (***Woodward et al., 2018***). In all optimizations, our STPT template was the fixed (target) image. This integration is facilitated by the fact that all current templates adopt the parcellation proposed by ***Paxinos et al***. (***2012***), ensuring uniformity of histological criteria and nomenclature across studies.

The mapping between the BMCA and BMA was done by computing the warping field between the STPT image template and the T2 weighted population average MRI template (isotropic voxel resolution of 100*μm*) of the BMA using ANTs.

The reference image of the MBCA was a 3D image stack of 63 cortical NISSL stained marmoset brain sections (825 × 63 × 550 voxels with a spatial resolution of 0.04 × 0.5 × 0.04 *μm*^3^). Compared to our STPT template, the sagittal resolution was rather low. To improve the registration, we added a mask for the cortex for both templates STPT and MBCA as an additional data term for the ANTs optimization (mean square error as metric). The cortex mask for the MBCA template was generated by fusing all cortical labels in the MBCA atlas.

The mapping between the BMCA and the MBM atlas was performed in two steps. The affine registration was done between the STPT template and the MBM symmetric T2 weighted image with 80*μm* resolution using normalized mutual information as a metric. We added the mean square distance between cortex masks in the SyN step.

Using the warp fields, we mapped the gray matter atlas labels of the BMA, the cortical labels of the MBCA atlas, and the cortical, subcortical and white matter labels of the MBM version 2 to the BMCA.

#### Flatmap-stack mapping

We mapped all 3D tracer image data in the cortex from the BMCA template image to 3D flatmap stacks. A flatmap stack is a 3D image representation of the cortex, where the XY-plane defines the position on the cortex surface and the z-direction defines the relative cortical depth. Flatmap stack mappings are extensions of the flatmaps which are part of the MBM atlas and the BMA atlas. Figure 12 shows an example of cortical anterograde tracer densities mapped to a flatmap stack.

Both the MBM atlas and the BMA atlas share triangulated 3D surfaces that map 3D points of the mid-surface of one hemisphere in the marmoset cortex to a 2D flatmap. The data were publicly available (MBM^1^ and BMA^2^). We exploited the data to map the entire cortex to a 3D image stack that extends the flatmaps with cortical depth.

We first used our warping fields to map the vertices of the 3D surfaces to our STPT template space. Then we defined the inner border and outer border of the cortex in the STPT template. This step was done manually using the image annotation function in the 3D Slicer tool. We computed the normals of the surface for the cortical surface, where the normals at the inside pointed towards the cortex, and the normals at the outside pointed away from the cortex. We then used heat propagation to diffuse the directional information within the entire cortex and normalize the result. The directional field defines trajectories that start at the inner cortex boundary, follow the directional field, and terminate at the outer cortex boundary.

We determined the trajectories that intersected with the vertices of the 3D mid-surface of the flatmap data. The 2D counterparts of the intersected flatmap vertices defined the flatmap stack XY coordinates. We traced the trajectories, and the relative position on the trajectory represented the stack depth.

Based on the stack of 3D image coordinates, we generated ANTS warping fields for both the MBM based flatmap stack and the BMA based flatmap stack. The target size of a flatmap stack was 500×500×50 voxels. The cortical atlas data of the BMA, MBCA, and MBM atlases have been mapped to the flatmap stack image space as well.

#### Diffusion MRI data

The individual HARDI data is also registered to our template space to construct a high-quality structural population average. Registration was done by ANTS using mutual information of the STPT template and T2/b0 contrast. Gradient orientations were re-oriented accordingly. After mapping, the original 128 directions were mapped to left-right symmetric 64 directions using spherical interpolation, and a left-right symmetric population average HARDI was generated. The final resolution of the average HARDI template is 200*μm*. From the template, we generated standard diffusion metrics such as diffusivities and fractional anisotropy. We further provide streamlines (fiber tracts) generated by global tractography (***Reisert et al., 2011***) and fiber track-density images (***Calamante et al., 2010***) for each injection site. Figure 13 outlines the generation of fiber track-density images.

**Figure 13.**
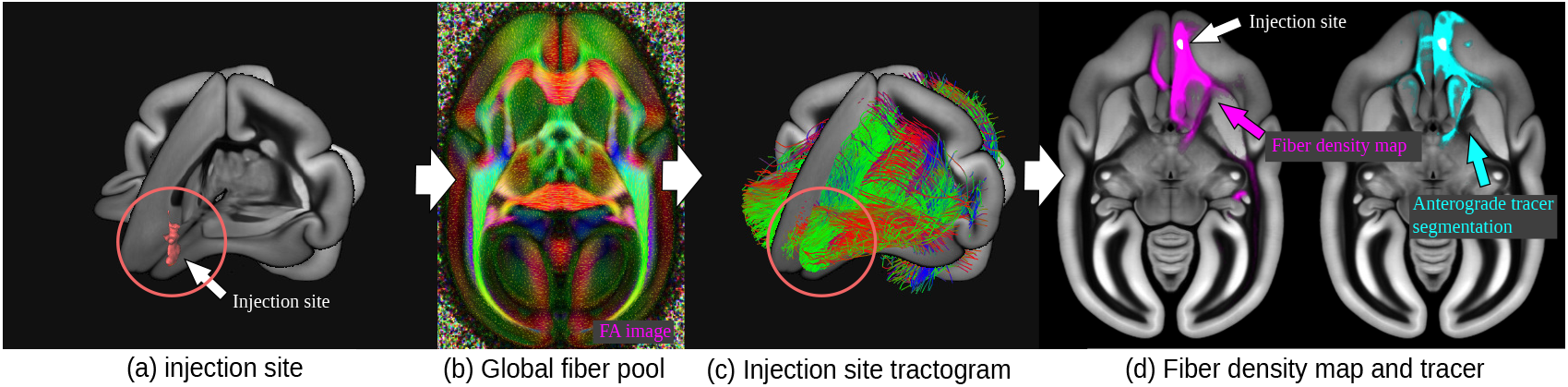
Generation of fiber track-density images: (a) based on an injection site location, (c,b) the pipeline selects all intersecting fibers from a global fiber pool (a large tractogram). (d) shows fiber density and tracer density, both associated with the same injection site.

#### Nissl and backlit

After the STPT imaging, the sections were imaged twice. Once before (backlit) and after Nissl staining. For backlit imaging, which reveals features of the brain myelination, the slices were collected from the STPT microscope and mounted onto slides. Then after imaging, stained for Nissl bodies and imaged a second time. In both steps, physical deformations happened due to the mounting, staining, or decaying processes. The savior deformations happened during the initial mount. We used a multi-modal image registration to undo the deformations in the images. Figure 14 outlines the registration pipeline. First, the Nissl image was mapped back to the backlit image. Then the backlit image was mapped back to its corresponding STPT image slice. Applying the concatenated warp fields mapped both images back to the image space of the original STPT image section. In an initial trial, the registration between the three different image modalities occasionally failed due to the major visual differences which hindered automation. We deployed a semi-supervised image-to-image translation that changed the contrast for the Nissl and backlit image to lookalike the STPT template image during the initial affine registration, which solved the problem. The image-to-image translation was realized with a generative adversarial network. Further details and code are publicly available online (***Skibbe et al., 2021***).

**Figure 14.**
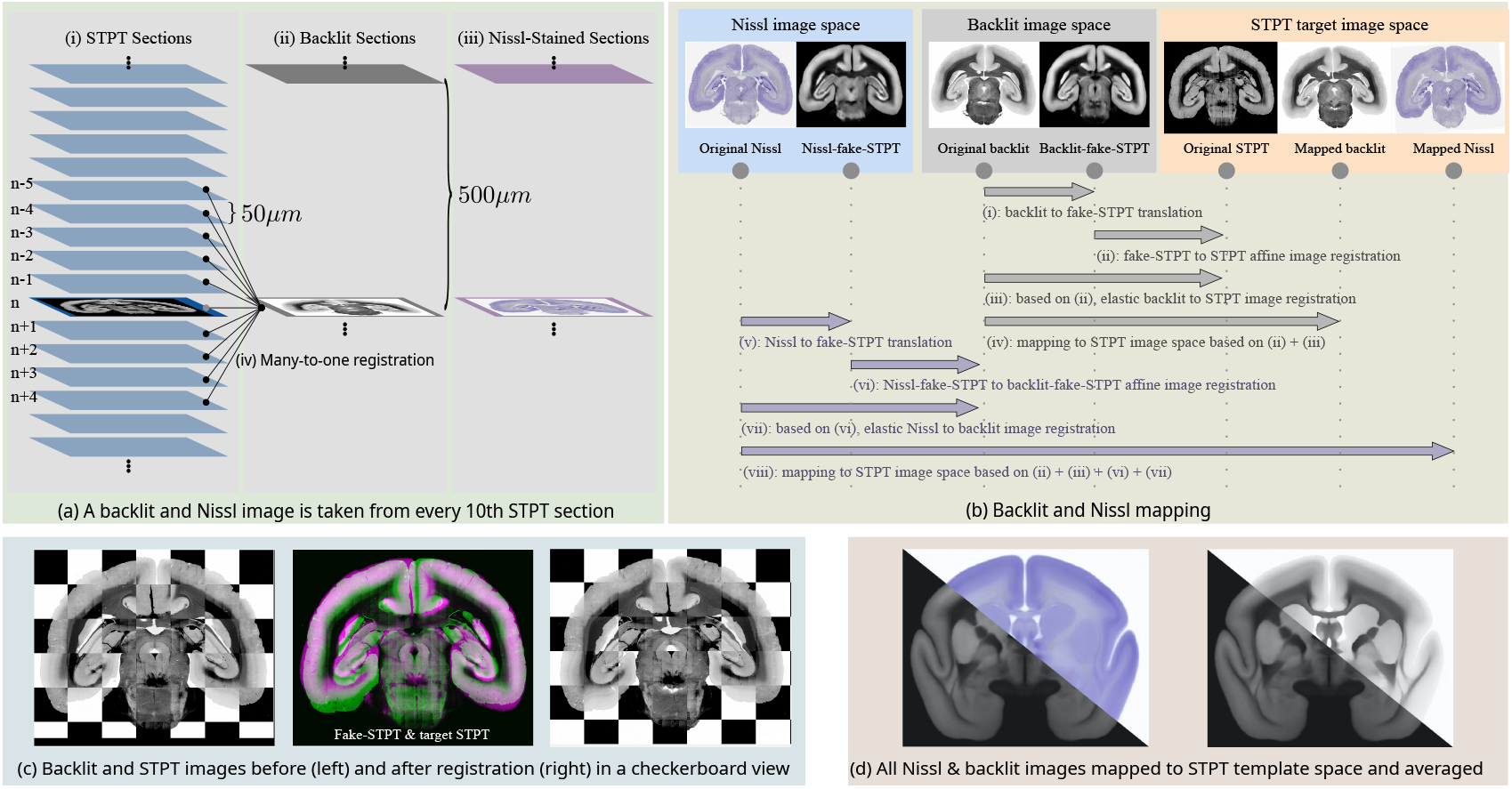
The Nissl and backlit registration pipeline. (a) For every 10th STPT tissue section, we take an additional backlit and a Nissl image with a second microscope. (b) Each time a section is moved or stained, it undergoes physical deformations. We established a robust image registration pipeline that reliably aligns the Backlit/Nissl images with the STPT images in a fully automated manner. (c) An example of a backlit image section before and after alignment. (d) After mapping the Nissl/Backlit images of our entire population to the STPT reference template, we computed population averages.

### Integration of the Marmoset Brain Connectivity Atlas

We used the Marmoset Brain Connectivity Atlas API^3^ (***Majka et al., 2020, 2016***) to download the retrograde cell data sets. We first got a list of all available datasets via the “injections” command. Then, for each injection, we downloaded the list of cell locations via the “cells?injection_id=“ command. The cell locations were given within the Paxinos stereotaxic reference space (***Paxinos et al., 2012***). Cells were labeled as “supragranular” and “infragranular” based on their cortical location with respect to cortical layer IV.

We used our ANTs transformation field to map the cell positions and injection site locations to the BMCA. For all injections, we created cell density images. We generated images with an isotropic spatial resolution of 100*μm*, and images with 400*μm* resolution. Similar to the data that is publicly available on the MBCA web portal, we generated three types of cell density maps. Each one for “infragranular”, “supragranular” and for combination of both cell categories. In addition, all density maps have been mapped to flatmap stacks using our ANTs transformation field.

### A diffeomorphic warp between marmoset and human brain

To map marmoset to human brain anatomy is a challenging task. In particular, the absence of cortical foldings in the marmoset does not allow image feature-based registrations based on the cortex. Hence, we concentrated on white matter anatomy. Diffusion MRI is particularly rich in structure in white matter areas. We used a mixture of manual and automatic coregistration steps. To represent human anatomy the MNI2009b symmetric template space was used. We constructed a diffusion MRI template in this space based on 200 HCP subjects (http://www.humanconnectomeproject.org). In a first step, rough correspondences were established by a downscaling of the human brain by a factor of 5 and using the manual deformation tool which comes as a part of NORA (https://www.noraimaging.com/) in our Nora-StackApp (see below). Based on the initial alignment, the ANTs coregistration toolbox was used to make a joint registration of directional diffusion MRI information and manually annotated contrasts (masks of the unique landmarks, like the anterior commissure, the subthalamic nucleus, or the corpus callosum). One particular challenge was the quite different ventricular anatomy of marmoset and human anatomy, which was not perfectly solvable by an automatic registration approach. So, the automatic registration was intervened by manual refinement steps guided by an experienced neuroscientist with neuroradiological background and solid knowledge of marmoset brain anatomy. Note that our whole approach is purely driven by anatomical features and does not incorporate any functional information.

#### White matter atlas comparison

From 28 regions in the JHU atlas (***Hua et al., 2008***; ***Wakana et al., 2007***; ***Mori et al., 2005***) (we merged regions from the left and right hemispheres), we selected 14 regions with similar ontology and shape in humans and marmosets (***Liu et al., 2020***). Corresponding labels are listed in table 1.

**Table 1.** The 14 regions in the JHU and MBM white matter atlases that we used for evaluating the warp between the marmoset brain and the human brain.

| JHU | MBM v2 (white matter atlas) |
| --- | --- |
| Anterior corona radiata | corona radiata |
| Medial lemniscus | medial lemniscus |
| Superior cerebellar peduncle | superior cerebellar peduncle |
| Inferior cerebellar peduncle | inferior cerebellar peduncle |
| Middle cerebellar peduncle | middle cerebellar peduncle |
| Uncinate fasciculus | uncinate fasciculus |
| Fornix (column and body of fornix) | body of fornix |
| Superior longitudinal fasciculus | superior longitudinal fasciculus |
| External capsule | external capsule |
| Cerebral peduncle | cerebral peduncle |
| Posterior limb of internal capsule | posterior limb of internal capsule |
| Anterior limb of internal capsule | anterior limb of internal capsule |
| Tapetum | tapetum |
| Body of corpus callosum | corpus callosum |

## Data availability

The **BMCA-Explorer** is publicly available under the following link https://bia.riken.jp/doku.php?id=tools:bmca. Login: “BMCA”, Password “BMCAACMB”. The **Nora-StackApp**, the STPT template, the HARDI population average template, the warping fields between various marmoset brain atlases, the flat map mapping, and the mapped data from the Marmoset Brain Connectivity Atlas are publicly available in the RIKEN CBS data repository https://neurodata.riken.jp/mdrs/explorer/?id=39&path=%2F (Password “BMCAACMB”).

The git repositories containing the source code for the pipeline and the code for generating the flatmap stack warping fields are publicly available here https://bitbucket.org/skibbe/tc_pipeline and here https://bitbucket.org/skibbe/flatmap_stack/, respectively.

The 3D image stacks of anterograde tracer will be publicly accessible at the RIKEN CBS data repository (https://neurodata.riken.jp/mdrs/explorer/?id=39&path=%2F) and the Brain/MINDS data portal after acceptance of this manuscript, and after acceptance of the paper by ***Watakabe et al***. (***2021***), which describes the anterograde core data.

## Acknowledgements

This work was supported by the program for Brain Mapping by Integrated Neurotechnologies for Disease Studies (Brain/MINDS) from the Japan Agency for Medical Research and Development AMED (JP15dm0207001). It was also supported by Scientific Research on Innovative Areas (22123009) from MEXT, Japan and National Science Centre of Poland (2019/35/D/NZ4/03031).

---

**Appendix 0 Figure 15.**
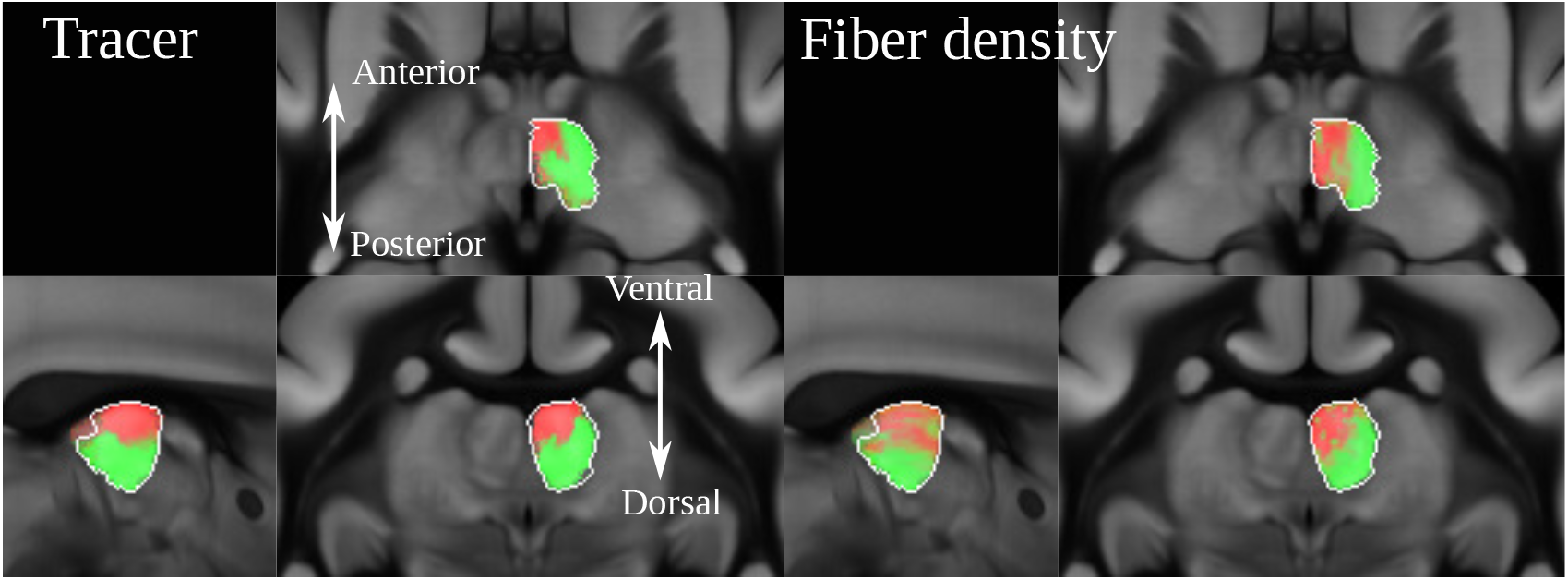
The image shows the strong projections of signals derived from anterograde tracing and dMRI in the mediodorsal thalamic nucleus. Both dMRI and anterograde tracer suggest that A32 projects anteromedially while A8aV projects posterolaterally.

**Appendix 0 Figure 16.**
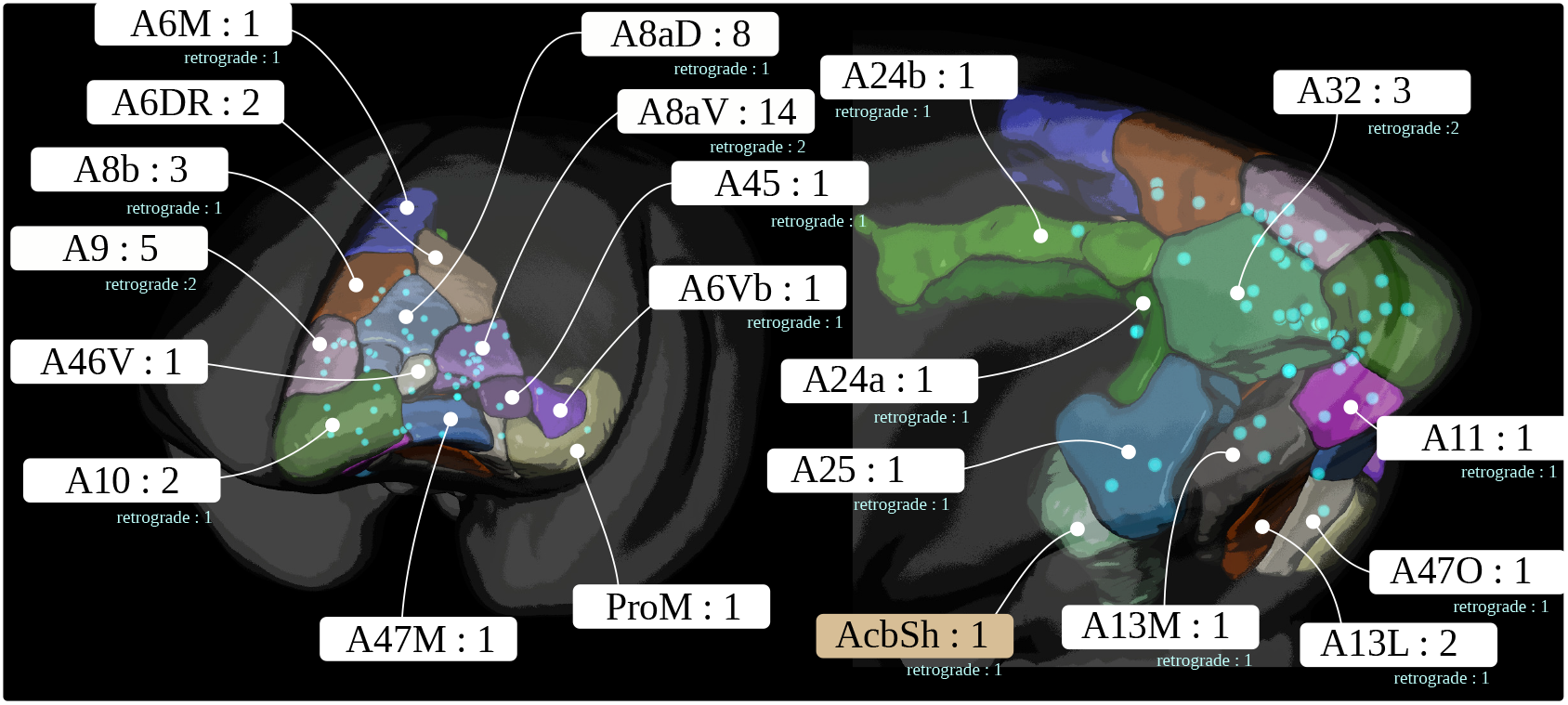
The figure shows the locations of all 52 anterograde, and corresponding 19 retrograde tracer injections in the marmoset prefrontal cortex.

## Footnotes

1 https://marmosetbrainmapping.org/atlas.html#v3

2 https://dataportal.brainminds.jp/atlas-package-download-main-page/bma-2019-ex-vivo

3 https://github.com/Neuroinflab/analysis.marmosetbrain.org/wiki/Application-Programming-Interface

## References

1. Ando K, Obayashi S, Nagai Y, Oh-Nishi A, Minamimoto T, Higuchi M, Inoue T, Itoh T, Suhara T. PET analysis of dopaminergic neurodegeneration in relation to immobility in the MPTP-treated common marmoset, a model for Parkinson’s disease. PLoS One. 2012;.

2. Avants BB, Tustison NJ, Song G, Cook PA, Klein A, Gee JC. A reproducible evaluation of ANTs similarity metric performance in brain image registration. Neuroimage. 2011; 54(3):2033–2044.

3. Bakker R, Wachtler T, Diesmann M. CoCoMac 2.0 and the future of tract-tracing databases. Frontiers in neuroinformatics. 2012; 6:30.

4. Bakola S, Burman KJ, Bednarek S, Chan JM, Jermakow N, Worthy KH, Majka P, Rosa MG. Afferent connections of cytoarchitectural area 6M and surrounding cortex in the marmoset: putative homologues of the supple-mentary and pre-supplementary motor areas. Cerebral Cortex. 2022; 32(1):41–62.

5. Belmonte JCI, Callaway EM, Caddick SJ, Churchland P, Feng G, Homanics GE, Lee KF, Leopold DA, Miller CT, Mitchell JF, et al. Brains, genes, and primates. Neuron. 2015; 86(3):617–631.

6. Burman KJ, Rosa MG. Architectural subdivisions of medial and orbital frontal cortices in the marmoset monkey (Callithrix jacchus). Journal of Comparative Neurology. 2009; 514(1):11–29.

7. Calamante F, Tournier JD, Jackson GD, Connelly A. Track-density imaging (TDI): super-resolution white matter imaging using whole-brain track-density mapping. Neuroimage. 2010; 53(4):1233–1243.

8. Carlén M. What constitutes the prefrontal cortex? Science. 2017; 358(6362):478–482.

9. Chaplin TA, Yu HH, Soares JG, Gattass R, Rosa MG. A conserved pattern of differential expansion of cortical areas in simian primates. Journal of Neuroscience. 2013; 33(38):15120–15125.

10. Felleman DJ, Van Essen DC. Distributed hierarchical processing in the primate cerebral cortex. Cerebral cortex (New York, NY: 1991). 1991; 1(1):1–47.

11. Girard G, Caminiti R, Battaglia-Mayer A, St-Onge E, Ambrosen KS, Eskildsen SF, Krug K, Dyrby TB, Descoteaux M, Thiran JP, et al. On the cortical connectivity in the macaque brain: A comparison of diffusion tractography and histological tracing data. Neuroimage. 2020; 221:117201.

12. Glasser MF, Coalson TS, Robinson EC, Hacker CD, Harwell J, Yacoub E, Ugurbil K, Andersson J, Beckmann CF, Jenkinson M, et al. A multi-modal parcellation of human cerebral cortex. Nature. 2016; 536(7615):171–178.

13. Goulas A, Majka P, Rosa MG, Hilgetag CC. A blueprint of mammalian cortical connectomes. PLoS biology. 2019; 17(3):e2005346.

14. Gutierrez CE, Skibbe H, Nakae K, Tsukada H, Lienard J, Watakabe A, Hata J, Reisert M, Woodward A, Yamaguchi Y, et al. Optimization and validation of diffusion MRI-based fiber tracking with neural tracer data as a reference. Scientific reports. 2020; 10(1):1–18.

15. Hioki H, Kuramoto E, Konno M, Kameda H, Takahashi Y, Nakano T, Nakamura KC, Kaneko T. High-level trans-gene expression in neurons by lentivirus with Tet-Off system. Neuroscience research. 2009; 63(2):149–154.

16. Hori Y, Cléry JC, Selvanayagam J, Schaeffer DJ, Johnston KD, Menon RS, Everling S. Interspecies activation correlations reveal functional correspondences between marmoset and human brain areas. Proceedings of the National Academy of Sciences. 2021; 118(37).

17. Hua K, Zhang J, Wakana S, Jiang H, Li X, Reich DS, Calabresi PA, Pekar JJ, van Zijl PC, Mori S. Tract probability maps in stereotaxic spaces: analyses of white matter anatomy and tract-specific quantification. Neuroimage. 2008; 39(1):336–347.

18. Ioffe S, Szegedy C. Batch normalization: Accelerating deep network training by reducing internal covariate shift. arXiv preprint 150203167. 2015;.

19. Jiang H, Van Zijl PC, Kim J, Pearlson GD, Mori S. DtiStudio: resource program for diffusion tensor computation and fiber bundle tracking. Computer methods and programs in biomedicine. 2006; 81(2):106–116.

20. Kita Y, Nishibe H, Wang Y, Hashikawa T, Kikuchi SS, Mami U, Yoshida AC, Yoshida C, Kawase T, Ishii S, et al. Cellular-resolution gene expression profiling in the neonatal marmoset brain reveals dynamic species-and region-specific differences. Proceedings of the National Academy of Sciences. 2021; 118(18).

21. Kuan L, Li Y, Lau C, Feng D, Bernard A, Sunkin SM, Zeng H, Dang C, Hawrylycz M, Ng L. Neuroinformatics of the allen mouse brain connectivity atlas. Methods. 2015; 73:4–17.

22. Liu C, Frank QY, Newman JD, Szczupak D, Tian X, Yen CCC, Majka P, Glen D, Rosa MG, Leopold DA, et al. A resource for the detailed 3D mapping of white matter pathways in the marmoset brain. Nature neuroscience. 2020; 23(2):271–280.

23. Liu C, Frank QY, Yen CCC, Newman JD, Glen D, Leopold DA, Silva AC. A digital 3D atlas of the marmoset brain based on multi-modal MRI. Neuroimage. 2018; 169:106–116.

24. Liu C, Yen CCC, Szczupak D, Tian X, Glen D, Silva AC. Marmoset Brain Mapping V3: Population multi-modal standard volumetric and surface-based templates. NeuroImage. 2021; 226:117620.

25. Maier-Hein KH, Neher PF, Houde JC, Côté MA, Garyfallidis E, Zhong J, Chamberland M, Yeh FC, Lin YC, Ji Q, et al. The challenge of mapping the human connectome based on diffusion tractography. Nature communications. 2017; 8(1):1–13.

26. Majka P, Bai S, Bakola S, Bednarek S, Chan JM, Jermakow N, Passarelli L, Reser DH, Theodoni P, Worthy KH, et al. Open access resource for cellular-resolution analyses of corticocortical connectivity in the marmoset monkey. Nature communications. 2020; 11(1):1–14.

27. Majka P, Bednarek S, Chan JM, Jermakow N, Liu C, Saworska G, Worthy KH, Silva AC, Wójcik DK, Rosa MG. Histology-Based Average Template of the Marmoset Cortex With Probabilistic Localization of Cytoarchitectural Areas. Neuroimage. 2021; 226:117625.

28. Majka P, Chaplin TA, Yu HH, Tolpygo A, Mitra PP, Wójcik DK, Rosa MG. Towards a comprehensive atlas of cortical connections in a primate brain: Mapping tracer injection studies of the common marmoset into a reference digital template. Journal of Comparative Neurology. 2016; 524(11):2161–2181.

29. Marcus D, Harwell J, Olsen T, Hodge M, Glasser M, Prior F, Jenkinson M, Laumann T, Curtiss S, Van Essen D. Informatics and data mining tools and strategies for the human connectome project. Frontiers in neuroinformatics. 2011; 5:4.

30. Markov NT, Ercsey-Ravasz MM, Ribeiro Gomes A, Lamy C, Magrou L, Vezoli J, Misery P, Falchier A, Quilodran R, Gariel MA, et al. A weighted and directed interareal connectivity matrix for macaque cerebral cortex. Cerebral cortex. 2014; 24(1):17–36.

31. Markov NT, Vezoli J, Chameau P, Falchier A, Quilodran R, Huissoud C, Lamy C, Misery P, Giroud P, Ullman S, et al. Anatomy of hierarchy: feedforward and feedback pathways in macaque visual cortex. Journal of Comparative Neurology. 2014; 522(1):225–259.

32. Milham M, Petkov C, Belin P, Hamed SB, Evrard H, Fair D, Fox A, Froudist-Walsh S, Hayashi T, Kastner S, et al. Toward next-generation primate neuroscience: A collaboration-based strategic plan for integrative neuroimaging. Neuron. 2022; 110(1):16–20.

33. Miller CT, Freiwald WA, Leopold DA, Mitchell JF, Silva AC, Wang X. Marmosets: a neuroscientific model of human social behavior. Neuron. 2016; 90(2):219–233.

34. Mori S, Wakana S, Van Zijl PC, Nagae-Poetscher L. MRI atlas of human white matter. Elsevier; 2005.

35. Oh SW, Harris JA, Ng L, Winslow B, Cain N, Mihalas S, Wang Q, Lau C, Kuan L, Henry AM, et al. A mesoscale connectome of the mouse brain. Nature. 2014; 508(7495):207–214.

36. Okano H, Mitra P. Brain-mapping projects using the common marmoset. Neuroscience research. 2015; 93:3–7.

37. Okano H, Miyawaki A, Kasai K. Brain/MINDS: brain-mapping project in Japan. Phil Trans R Soc B. 2015; 370(1668):20140310.

38. Okano H, Sasaki E, Yamamori T, Iriki A, Shimogori T, Yamaguchi Y, Kasai K, Miyawaki A. Brain/MINDS: A Japanese National Brain Project for Marmoset Neuroscience. Neuron. 2016; 92(3):582–590.

39. Okano H. Current status of and perspectives on the application of marmosets in neurobiology. Annual Review of Neuroscience. 2021; 44:27–48.

40. Paxinos G, Watson C, Petrides M, Rosa M, Tokuno H. The marmoset brain in stereotaxic coordinates. Elsevier Academic Press; 2012.

41. Phillips JM, Fish LR, Kambi NA, Redinbaugh MJ, Mohanta S, Kecskemeti SR, Saalmann YB. Topographic organization of connections between prefrontal cortex and mediodorsal thalamus: Evidence for a general principle of indirect thalamic pathways between directly connected cortical areas. Neuroimage. 2019; 189:832–846.

42. Ragan T, Kadiri LR, Venkataraju KU, Bahlmann K, Sutin J, Taranda J, Arganda-Carreras I, Kim Y, Seung HS, Osten P. Serial two-photon tomography for automated ex vivo mouse brain imaging. Nature methods. 2012; 9(3):255–258.

43. Reisert M, Mader I, Anastasopoulos C, Weigel M, Schnell S, Kiselev V. Global fiber reconstruction becomes practical. Neuroimage. 2011; 54(2):955–962.

44. Reser DH, Burman KJ, Yu HH, Chaplin TA, Richardson KE, Worthy KH, Rosa MG. Contrasting patterns of cortical input to architectural subdivisions of the area 8 complex: a retrograde tracing study in marmoset monkeys. Cerebral Cortex. 2013; 23(8):1901–1922.

45. Rockland KS. About connections. Frontiers in Neuroanatomy. 2015; 9. https://www.frontiersin.org/article/10.3389/fnana.2015.00061, doi: 10.3389/fnana.2015.00061.

46. Ronneberger O, Fischer P, Brox T. U-net: Convolutional networks for biomedical image segmentation. In: International Conference on Medical image computing and computer-assisted intervention Springer; 2015. p. 234–241.

47. Rosa MG, Soares JG, Chaplin TA, Majka P, Bakola S, Phillips KA, Reser DH, Gattass R. Cortical afferents of area 10 in Cebus monkeys: implications for the evolution of the frontal pole. Cerebral Cortex. 2019; 29(4):1473–1495.

48. Sadakane O, Masamizu Y, Watakabe A, Terada SI, Ohtsuka M, Takaji M, Mizukami H, Ozawa K, Kawasaki H, Matsuzaki M, et al. Long-term two-photon calcium imaging of neuronal populations with subcellular resolution in adult non-human primates. Cell reports. 2015; 13(9):1989–1999.

49. Sato K, Sasaguri H, Kumita W, Inoue T, Kurosaki Y, Nagata K, Mihira N, Sato K, Sakuma T, Yamamoto T, et al. A non-human primate model of familial Alzheimer’s disease. bioRxiv. 2020;.

50. Schaeffer DJ, Hori Y, Gilbert KM, Gati JS, Menon RS, Everling S. Divergence of rodent and primate medial frontal cortex functional connectivity. Proceedings of the National Academy of Sciences. 2020; 117(35):21681–21689.

51. Skibbe H, Watakabe A, Nakae K, Gutierrez CE, Tsukada H, Hata J, Kawase T, Gong R, Woodward A, Doya K, et al. MarmoNet: a pipeline for automated projection mapping of the common marmoset brain from whole-brain serial two-photon tomography. arXiv preprint 190800876. 2019;.

52. Skibbe H, Watakabe A, Rachmadi F, Gutierrez CE, Nakae K, Yamamori T. Semi-supervised Image-to-Image translation for robust image registration. In: Medical Imaging with Deep Learning (MIDL); 2021..

53. Sneve MH, Grydeland H, Rosa MG, Paus T, Chaplin T, Walhovd K, Fjell AM. High-expanding regions in primate cortical brain evolution support supramodal cognitive flexibility. Cerebral Cortex. 2019; 29(9):3891–3901.

54. Solomon SG, Rosa MG. A simpler primate brain: the visual system of the marmoset monkey. Frontiers in neural circuits. 2014; 8:96.

55. Srivastava N, Hinton G, Krizhevsky A, Sutskever I, Salakhutdinov R. Dropout: a simple way to prevent neural networks from overfitting. The journal of machine learning research. 2014; 15(1):1929–1958.

56. Stephan KE, Kamper L, Bozkurt A, Burns GA, Young MP, Kötter R. Advanced database methodology for the Collation of Connectivity data on the Macaque brain (CoCoMac). Philosophical Transactions of the Royal Society of London Series B: Biological Sciences. 2001; 356(1412):1159–1186.

57. Theodoni P, Majka P, Reser DH, Wójcik DK, Rosa MG, Wang XJ. Structural attributes and principles of the neocortical connectome in the marmoset monkey. Cerebral Cortex. 2022; 32(1):15–28.

58. Thomas C, Frank QY, Irfanoglu MO, Modi P, Saleem KS, Leopold DA, Pierpaoli C. Anatomical accuracy of brain connections derived from diffusion MRI tractography is inherently limited. Proceedings of the National Academy of Sciences. 2014; 111(46):16574–16579.

59. Toarmino CR, Yen CC, Papoti D, Bock NA, Leopold DA, Miller CT, Silva AC. Functional magnetic resonance imaging of auditory cortical fields in awake marmosets. Neuroimage. 2017; 162:86–92.

60. Tournier JD, Calamante F, Connelly A. MRtrix: diffusion tractography in crossing fiber regions. International journal of imaging systems and technology. 2012; 22(1):53–66.

61. Wakana S, Caprihan A, Panzenboeck MM, Fallon JH, Perry M, Gollub RL, Hua K, Zhang J, Jiang H, Dubey P, et al. Reproducibility of quantitative tractography methods applied to cerebral white matter. Neuroimage. 2007; 36(3):630–644.

62. Wang XJ, Pereira U, Rosa MG, Kennedy H. Brain connectomes come of age. Current Opinion in Neurobiology. 2020; 65:152–161.

63. Watakabe A, Skibbe H, Nakae K, Abe H, Ichinohe N, Wang J, Takaji M, Mizukami H, Woodward A, Gong R, Hata J, Okano H, Ishii S, Yamamori T. Connectional architecture of the prefrontal cortex in the marmoset brain. bioRxiv. 2021; p. 2021.12.26.474213. doi: 10.1101/2021.12.26.474213.

64. Watakabe A, Takaji M, Kato S, Kobayashi K, Mizukami H, Ozawa K, Ohsawa S, Matsui R, Watanabe D, Yamamori T. Simultaneous visualization of extrinsic and intrinsic axon collaterals in Golgi-like detail for mouse corticothalamic and corticocortical cells: a double viral infection method. Frontiers in neural circuits. 2014; 8:110.

65. Watanabe S, Kurotani T, Oga T, Noguchi J, Isoda R, Nakagami A, Sakai K, Nakagaki K, Sumida K, Hoshino K, et al. Functional and molecular characterization of a non-human primate model of autism spectrum disorder shows similarity with the human disease. Nature communications. 2021; 12(1):1–13.

66. Woodward A, Hashikawa T, Maeda M, Kaneko T, Hikishima K, Iriki A, Okano H, Yamaguchi Y. The Brain/MINDS 3D digital marmoset brain atlas. Scientific data. 2018; 5:180009.

67. Zingg B, Hintiryan H, Gou L, Song MY, Bay M, Bienkowski MS, Foster NN, Yamashita S, Bowman I, Toga AW, et al. Neural networks of the mouse neocortex. Cell. 2014; 156(5):1096–1111.

